# Joint actions of diverse transcription factor families establish neuron-type identities and promote enhancer selectivity

**DOI:** 10.1101/2020.09.04.283036

**Authors:** Angela Jimeno-Martín, Erick Sousa, Rebeca Brocal-Ruiz, Noemi Daroqui, Miren Maicas, Nuria Flames

## Abstract

To systematically investigate the complexity of neuron-specification regulatory networks we performed an RNA interference (RNAi) screen against all 875 transcription factors (TFs) encoded in *Caenorhabditis elegans* genome and searched for defects in nine different neuron types of the monoaminergic (MA) superclass and two cholinergic motoneurons.

We identified 91 TF candidates to be required for correct generation of these neuron types of which 28 were confirmed by mutant analysis. We found that correct reporter expression in each individual neuron type requires at least nine different TFs. Individual neuron types do not usually share TFs involved in their specification but share a common pattern of TFs belonging to the five most common TF families: Homeodomain (HD), basic Helix Loop Helix (bHLH), Zinc Finger (ZF), Basic Leucine Zipper Domain (bZIP) and Nuclear Hormone Receptors (NHR). HD TF members are over-represented, supporting a key role for this family in the establishment of neuronal identities. These five TF families area also prevalent when considering mutant alleles with previously reported neuronal phenotypes in *C. elegans*, *Drosophila* or mouse. In addition, we studied terminal differentiation complexity focusing on the dopaminergic terminal regulatory program. We found two HD TFs (UNC-62 and VAB-3) that work together with known dopaminergic terminal selectors (AST-1, CEH-43, CEH-20). Combined TF binding sites for these five TFs constitute a cis-regulatory signature enriched in the regulatory regions of dopaminergic effector genes. Our results provide new insights on neuron-type regulatory programs in *C. elegans* that could help better understand neuron specification and evolution of neuron types.

## INTRODUCTION

Cell diversity is particularly extensive in nervous systems because the complexity of neural function demands a remarkable degree of cellular specialization. Transcription factors (TFs) are the main orchestrators of neurontype specification/differentiation programs and induced expression of small combinations of TFs is sufficient for direct reprogramming of non-neuronal cells into neuron-like cells (Masserdotti et al. 2016).

Previous studies have identified some conserved features of neuron specification and differentiation programs in different neuron types and organisms. For example, the importance of signal-regulated TFs that mediate morphogens and intercellular signaling for lineage commitment and neuronal progenitor patterning (Borello and Pierani 2010; Andrews et al. 2019; Liu and Niswander 2005; Angerer et al. 2011; Nuez and Félix 2012; Rentzsch et al. 2017), the central role of specific basic Helix Loop Helix (bHLH) TFs as proneural factors (Bertrand et al. 2002; Guillemot and Hassan 2017), the key role of homeodomain (HD) TFs in neuron subtype specification (Briscoe et al. 2000; Shirasaki and Pfaff 2002; Thor et al. 1999; Durham et al. 2021) or the terminal selector model for neuronal terminal differentiation, in which specific TFs, termed terminal selectors, directly co-regulate expression of most neurontype specific effector genes (Hobert, 2008).

However, the complete gene regulatory networks that implement specific neuron identities, either during development or induced by reprogramming, are still poorly understood. To increase our global knowledge on this process we took advantage of the amenability of *Caenorhabditis elegans* for unbiased large-scale screens and performed an RNA interference (RNAi) screen against all 875 TFs encoded by the *C. elegans* genome. We systematically assessed their contribution in correct reporter gene expression in nine different types of neurons of the monoaminergic (MA) superclass and two cholinergic motoneurons. We focused mainly on MA neurons not only because they are evolutionary conserved and clinically relevant in humans (Flames and Hobert 2011), but also because the MA superclass comprises a set of neuronal types with very diverse developmental origins and functions in both worms and humans (Flames and Hobert 2011). We have previously shown that gene regulatory networks directing the terminal fate of two types of MA neurons (dopaminergic and serotonergic neurons) are conserved in worms and mammals (Lloret-Fernández et al. 2018; Doitsidou et al. 2013; Flames and Hobert 2009; Remesal et al. 2020). Thus, the identification of common principles underlying MA specification could help unravel general rules for neuron-type specification.

## RESULTS

### Whole genome transcription factor RNAi screen identifies new TFs required for neuron-type specification

The TF RNAi screen was performed feeding *rrf-3*(*pk1426*) mutant strain, that sensitizes neurons for RNAi effects (Simmer et al. 2003), with 875 different RNAi clones targeting all *C. elegans* TFs (**Supplemental Table S1**). To assess for neuron specific defects, *rrf-3*(*pk1426*) mutation was combined with three different fluorescent reporters that label the MA system in the worm (**Fig. 1A**): the vesicular monoamine transporter *otIs224*(*cat-1p::gfp*), expressed in all MA neurons; the dopamine transporter, *otIs181*(*dat-1p::mcherry*), expressed in dopaminergic neurons; and the tryptophan hydroxylase enzyme *vsIs97*(*tph-1p::dsred*), expressed in serotonergic neurons. Altogether our strategy labels nine different MA neuronal classes (dopaminergic ADE, CEPV, CEPD and PDE; serotonergic NSM, ADF and HSN; octopaminergic RIC and tyraminergic RIM) and the two cholinergic VC4 and VC5 motoneurons, which are not MA but express *cat-1* reporter for unknown reasons. Two additional MA neurons, AIM and RIH are not labeled by *otIs224*(*cat-1p::gfp*) and thus were not considered in our study. Analyzed neurons are developmentally, molecularly and functionally very diverse: (1) they arise from different branches of the AB lineage (**Figure 1B**), (2) they include motoneurons, sensory neurons, and interneurons, that altogether use five different neurotransmitters (**Fig. 1A**) and (3) each neuronal type expresses different transcriptomes, having in common the genes related to MA metabolism (**Fig. 1C**). In summary, considering the diversity of labeled neurons, we reasoned that their global study could unravel shared principles of *C. elegans* neuron specification and differentiation.

**Figure 1.**
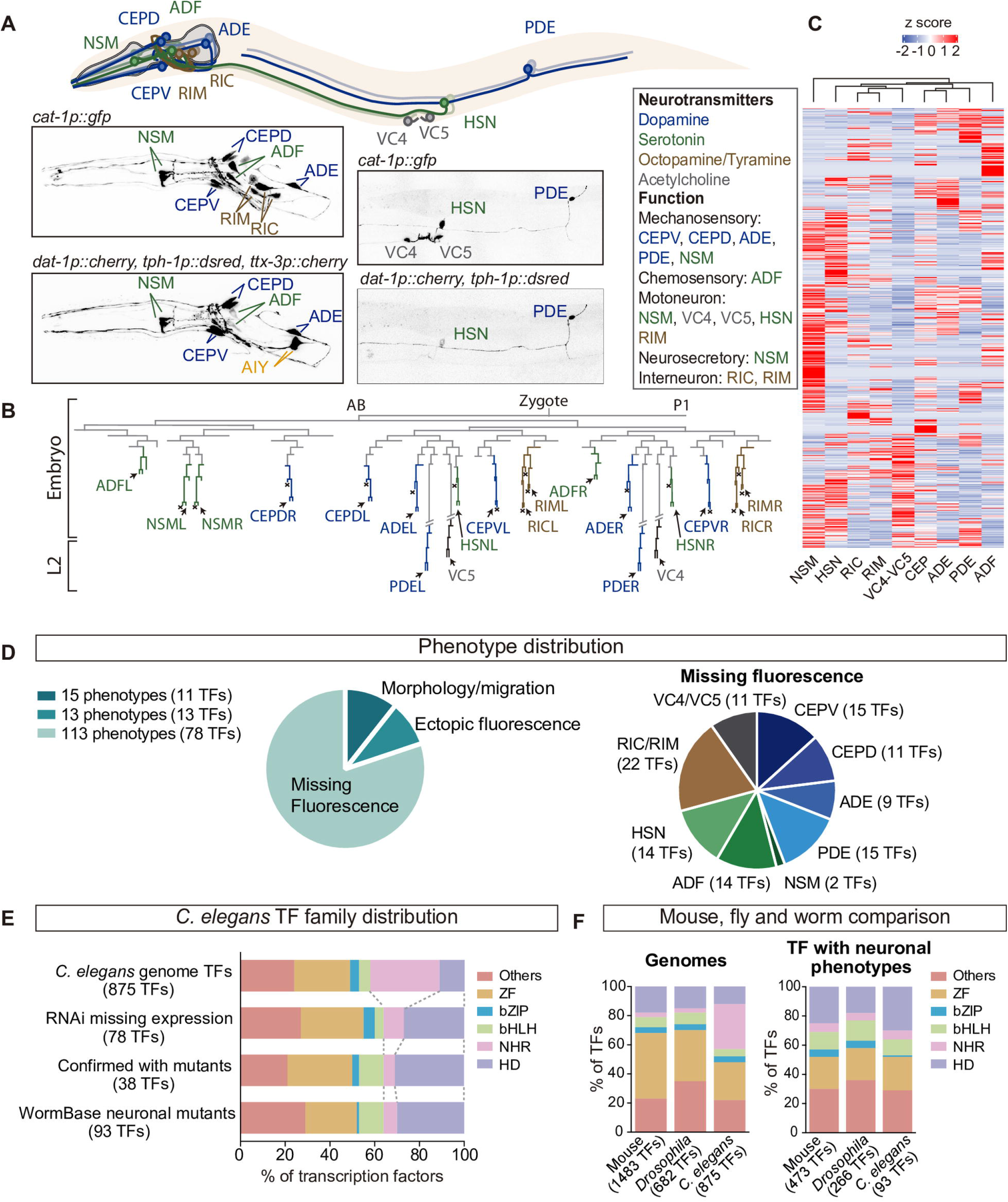
A genome wide transcription factor RNAi screen reveals specific TF families are required for the specification of eleven neuron types. **A**) *otIs224*(*cat-1p::gfp*) reporter strain labels nine classes of MA neurons (dopaminergic in blue, serotonergic in green and tyraminergic and octopaminergic in brown) and the VC4 and VC5 cholinergic motoneurons. AIM and RIH serotonergic neurons are not labeled by this reporter. HSN is both serotonergic and cholinergic. For the RNAi screen the dopaminergic *otIs181*(*dat-1p::mcherry*) and serotonergic *vsIs97*(*tph-1p::dsred*) reporters were also scored together with *otIs224. ttx-3p:mcherry* reporter, labeling AIY is cointegrated in *otIs181* but was not scored. **B**) Developmental *C. elegans* hermaphrodite lineage showing the diverse origins of neurons analyzed in this study. **C**) Heatmap showing the disparate transcriptomes of the different MA neurons. Data obtained from Larval L4 single-cell RNA-seq experiments (Taylor et al. 2021). **D**) Phenotype distribution of TF RNAi screen results. 91 TF RNAi clones produce 141 phenotypes as some TF RNAi clones are assigned to more than one cell type and/or phenotypic category. Most neuron types are affected by knock down of at least 9 different TFs. We could not differentiate between RIC and RIM due to proximity and morphological similarity; thus they were scored as a unique category. **E**) TF family distribution of TFs in *C. elegans* genome, TFs retrieved in our RNAi screen, TFs confirmed by mutant analysis and mutant alleles with any assigned neuronal phenotype in WormBase. Homeodomain TFs are over-represented and NHR TFs decreased compared to the genome distribution. **F**) Comparison of mouse, *Drosophila* and *C. elegans* genomic TF family distribution and of TFs with assigned neuronal phenotypes. Distribution of mouse and *Drosophila* families with phenotypes is more similar to *C. elegans* than genomic distributions. Moreover, mouse HD prevalence in neuronal phenotypes is increased compared to genomic HD frequency, similar to *C. elegans*.

From our screen, 91 of the 875 TF RNAi clones displayed a phenotype classified as either missing fluorescent cells, ectopic fluorescent cells, migration, axon guidance, morphology defects or combinations of them.

Considering each neuron type individually a total of 141 different phenotypes were identified, being missing or reduced reporter expression the most frequent (**Fig. 1D** and **Supplemental Table S2**).

We retrieved phenotypes for 24 out of 31 known regulators of MA neuron fate (**Supplemental Table S3**). Thus, we estimated a false negative rate of around 23%. To estimate the false positive rate of the screen we focused on the validation of the set of TFs displaying missing fluorescence phenotypes with penetrance higher than 20% (**Table 1** and **Supplemental Table S2**). 40 out of 46 RNAi phenotypes were verified by the corresponding mutant analysis (**Table 1** and **Supplemental Table S4**), estimating a false positive rate of 14%. See **Table 2** for a summary of RNAi screen hits, validations and error rates.

**Table 1.**
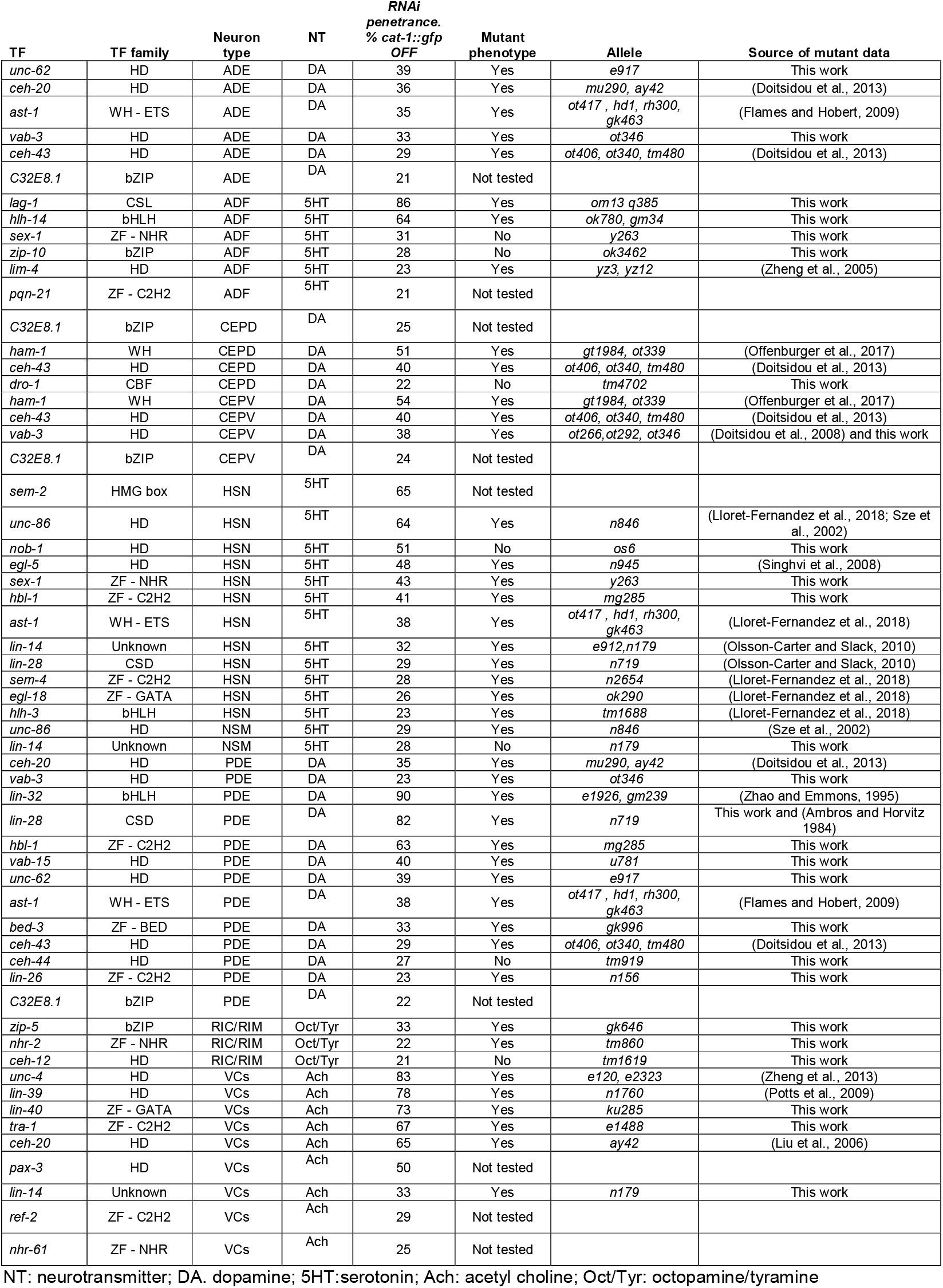
TF RNAi clones associated to missing reporter expression with > 20 % penetrance.

**Table 2.** RNAi screen summary.

| RNAi screen summary |  |
| --- | --- |
| # of RNAi phenotypes (all) | 141 |
| # of RNAi phenotypes (missing fluorescence) | 113 |
| # of RNAi phenotypes >20% penetrance (missing fluorescence) | 59 |
| # of TFs with RNAi phenotype (all) | 91 |
| # of TFs with RNAi phenotype (missing fluorescence) | 78 |
| # of TFs with RNAi phenotype >20% penetrance (missing fluorescence) | 39 |
| # of phenotypes confirmed by mutant alleles (missing fluorescence) | 52 |
| # of TFs with phenotypes confirmed by mutant alleles (missing fluorescence) | 38 |
| # of TFs with newly assigned neuronal phenotypes confirmed by mutant alleles (missing fluorescence) | 11 |
| False negative rate (TFs) | 23% |
| False positive rate (referred to phenotypes, >20% penetrance) | 14% |
| False positive rate (referred to TFs, >20% penetrance) | 15% |

Each neuron type was affected by 9 to 15 TF RNAi clones (**Fig. 1D** and **Supplemental Table S2**) with the exception of the NSM neurons, affected only by two clones producing missing fluorescence phenotypes. We noticed that NSM neurons showed weak phenotypes upon *gfp* RNAi treatment (**Supplemental Fig. S1**) implying that, even in the *rrf-3*(*pk1426*) sensitized background NSM could be particularly refractory to RNAi. Thus, we excluded NSM for further analysis. None of the TF RNAi clones affect all MA neurons, suggesting the absence of global regulators of MA effector gene expression or survival. Alternatively, global regulators of MA fate could show redundant functions or produce early embryonic lethality. Nevertheless, only 37 of the 875 TF RNAi clones produced embryonic lethality with no escapers. These clones were scored in the parental generation, six of them produced visible neuronal phenotypes and were included in further analyses (**Supplemental Table S2**). TFs with known roles in MA specification that were retrieved from the RNAi screen act at different developmental stages. For example, *lag-1* CSL TF is expressed in the postmitotic ADF neuron but not earlier, acting as terminal selector (Maicas et al. 2021), *lim-4* HD TF is expressed in the mother cell of ADF but not in the postmitotic neuron and regulates *lag-1* expression (Zheng et al. 2005; Maicas et al. 2021), while *hlh-14* bHLH TF is expressed earlier in the lineage and its proneural activity is required to generate the variety of neurons arising from that lineage (Masoudi et al. 2021; Poole et al. 2011). In addition, known TFs affecting PDE reporter expression also act at several steps of the specification/differentiation process: *ceh-16* HD TF is required for correct asymmetric divisions of the PDE lineage (V5 progenitor) (Huang et al. 2009), *lin-32* bHLH TF is a proneurogenic factor needed for the generation of PDE and its sister cell PVD neuron, similar to the role of *hlh-14* in ADF (Zhao and Emmons 1995) and *ast-1* ETS TF affects terminal fate of PDE but not PVD neuron (Flames and Hobert 2009). To expand on the analysis of different TFs acting on the same neuron type, we focused on novel PDE validated mutants. In *wild type* animals, panneuronal reporter *rab-3* GTPase is expressed both in PDE and PVD neurons. HD *unc-62*(*e917*) and *vab-15*(*u781*) mutants show defects in *rab-3* expression both in PDE and PVD neurons. PDE and PVD specific reporters (*cat-1* for PDE and *dop-3* or *unc-86* for PVD) are also affected in these mutants (**Supplemental Fig. S2**), suggesting either lineage specification or broad PDE/PVD differentiation defects. Zinc Finger (ZF) C2H2 *hbl-1*(*mg285*) mutants display *rab-3* reporter expression only in one of the two neurons. Missing *rab-3* reporter expression coincides with lack of *dop-3* PVD reporter, suggesting *hbl-1* mutants affect PVD neuron specification and/or generation more than PDE (**Supplemental Fig. S2**). Finally, the unknown TF type *lin-14*(*n179*) shows normal *rab-3* reporter expression in PDE and PVD but defects in neuron-type specific markers of both neuron types, supporting a role in terminal differentiation but not in neuronal-lineage commitment (**Supplemental Fig. S2**). Thus, newly identified PDE mutants show a variety of phenotypes likely reflecting different TF actions at particular developmental stages.

### A specific set of transcription factor families controls neuron specification

Next, we focused on the 113 missing reporter expression RNAi phenotypes for different neuron types (assigned to 78 TF RNAi clones) and analyzed the screen results based on TF families instead of individual TF members. According to their DNA-binding domain, *C. elegans* TFs can be classified into more than fifty different TF families (Stegmaier et al. 2004; Narasimhan et al. 2015) (**Supplemental Table S1**). bHLH, HD, ZF, Basic Leucine Zipper Domain (bZIP) and Nuclear Hormone Receptor (NHR) families comprise 75% of the TFs in the *C. elegans* genome. RNAi clones targeting these families also generate 75% of the missing reporter phenotypes (**Fig. 1E**). We noticed that the prevalence of two TF families, HD and NHR, largely differs from what would be expected from the number of TF members encoded in the genome (**Fig. 1E**). HD family represents 12% of total TFs in the *C. elegans* genome but accounted for the 27% (21/78) of TF RNAi clones showing a phenotype. Thus, HD family is significantly over-represented in our screening (p<0.005, Fisher’s exact test).

Conversely, NHR TFs are under-represented when considering the total number of NHR TFs encoded in the genome (30% of all *C. elegans* TFs vs 9% of NHR RNAi clones with phenotype, p<0.0001, Fisher’s exact test) (**Fig. 1E**). A roughly similar TF family distribution is observed when considering MA neuron types individually (**Supplemental Fig. S3**). As the specific TF family distribution found in our RNAi screen could be biased by our false positive and negative rates, we focused only on the set of 38 TFs confirmed by mutant analysis. Mutant-verified TFs showed a similar TF family distribution (30% HD TFs and 5% NHR TFs; significantly different from genome distribution p<0.0001 and p=0.0017 respectively, Fisher’s exact test), suggesting a prevalent role for HD family in neuron specification (**Fig. 1E**).

Next, to explore whether this TF family distribution could apply to other neuron types, we analyzed the TF family distribution of the 93 *C. elegans* TFs for which mutant alleles display any neuronal phenotype according to WormBase data (**Supplemental Table S5**). TF family distribution for these 93 TFs is highly similar to the observed in our RNAi screen and our mutant analysis (**Fig. 1E**), suggesting that HD over- and NHR under-representation could constitute a general rule of the regulatory networks controlling neuronal identity in *C. elegans*.

NHR TF family is expanded in *C. elegans* (composed of 272 members) compared human genome (less than 50 members). Only 8% of the 272 *C. elegans* NHRs have orthologs in non-nematode species (Maglich et al. 2001; Taubert et al. 2011; Bodofsky et al. 2017). We found that phylogenetically conserved NHRs are enriched for neuronal phenotypes both in the MA RNAi screen (3 out of 7 NHR TFs) and in the WormBase neuronal mutants (5 out of 6 NHR TFs). This observation suggests that, among NHR members, those phylogenetically conserved could have a prevalent role in neuron specification. Alternatively, lack of phenotypes for nematode specific NHRs could be attributed to more redundant actions among them or to specialized functions in particular neuron types and gene targets as recently suggested (Sural and Hobert 2021).

Finally, we expanded our analysis to *Drosophila* and mouse model systems. The same five *C. elegans* most preponderant TF families constitute the majority of TFs also in *Drosophila* and mouse genomes, however, the number of NHR TFs is considerably smaller than in *C. elegans* (**Fig. 1F** and **Supplemental Table S6**). TF family distribution of 266 *Drosophila* and 473 Mouse TFs whose mutations generate neuronal phenotypes (source FlyBase and Mouse Genome Informatics) is more similar to *C.elegans* neuronal mutant distribution than the genomic distributions. HD over-representation is also found for mouse TFs with assigned neuronal phenotypes (18% of TFs in the mouse genome *vs* 25% of TFs with neuronal phenotypes, p<0.0002 Fisher’s exact test) although HD enrichment is not observed in *Drosophila* (**Fig. 1F** and **Supplemental Table S6**).

Thus, we found a stereotypic TF family distribution associated to generation, specification, differentiation, survival and/or function of different neuronal types in different model organisms with high prevalence of HD TFs.

### Identification of new transcription factors involved in dopaminergic terminal differentiation

Next, we aimed to use our TF RNAi screen data to study the complexity of terminal differentiation programs. In the immature postmitotic neuron, terminal differentiation is regulated by specific TFs, termed terminal selectors, that directly activate the expression of effector genes (*e.g*. ion channels, neurotransmitter receptors or biosynthesis enzymes) that define the neurontype specific transcriptome (Hobert 2008). Terminal selectors, like any other TFs, act in combinations to activate target enhancers, but the complexity of these terminal selector codes is not known. In the HSN serotonergic neuron, a combination of six transcription factors act as a terminal selector collective to control terminal differentiation of this neuron type (Lloret-Fernández et al. 2018). Thus, we aimed to explore whether other neuronal terminal differentiation programs exhibited similar regulatory complexity.

Defects in early lineage specification, neuronal terminal differentiation or survival can produce similar reporter expression defects. Thus early and late functions of TFs cannot be discerned *a priori* in our RNAi screen. To circumvent this limitation, we decided to focus on the four dopaminergic neuron subtypes (CEPV, CEPD, ADE and PDE) that arise from different lineages but converge to the same terminal regulatory program (**Fig. 2A**). Known early lineage determinants for dopaminergic neurons affect unique subtypes, for example, the already mentioned role of CEH-16/HD TF in V5 lineage (Huang et al. 2009) controls PDE generation but not other dopaminergic subtypes (**Supplemental Table S2**). Conversely, known dopaminergic terminal selectors TFs (*ast-1* ETS, *ceh-43* HD and *ceh-20* HD) act in all four neuron subtypes (Doitsidou et al. 2013; Flames and Hobert 2009) (**Fig. 2A** and **Supplemental Fig. S4**). Accordingly, we reasoned that RNAi clones leading to similarly broad dopaminergic phenotypes constitute good candidates to play a role in dopaminergic terminal differentiation. Therefore, we focused on *unc-62/MEIS-* HD and *vab-3*/PAIRED-HD since these TFs showed high penetrant RNAi phenotypes affecting several dopaminergic subtypes (**Table 1**).

**Figure 2.**
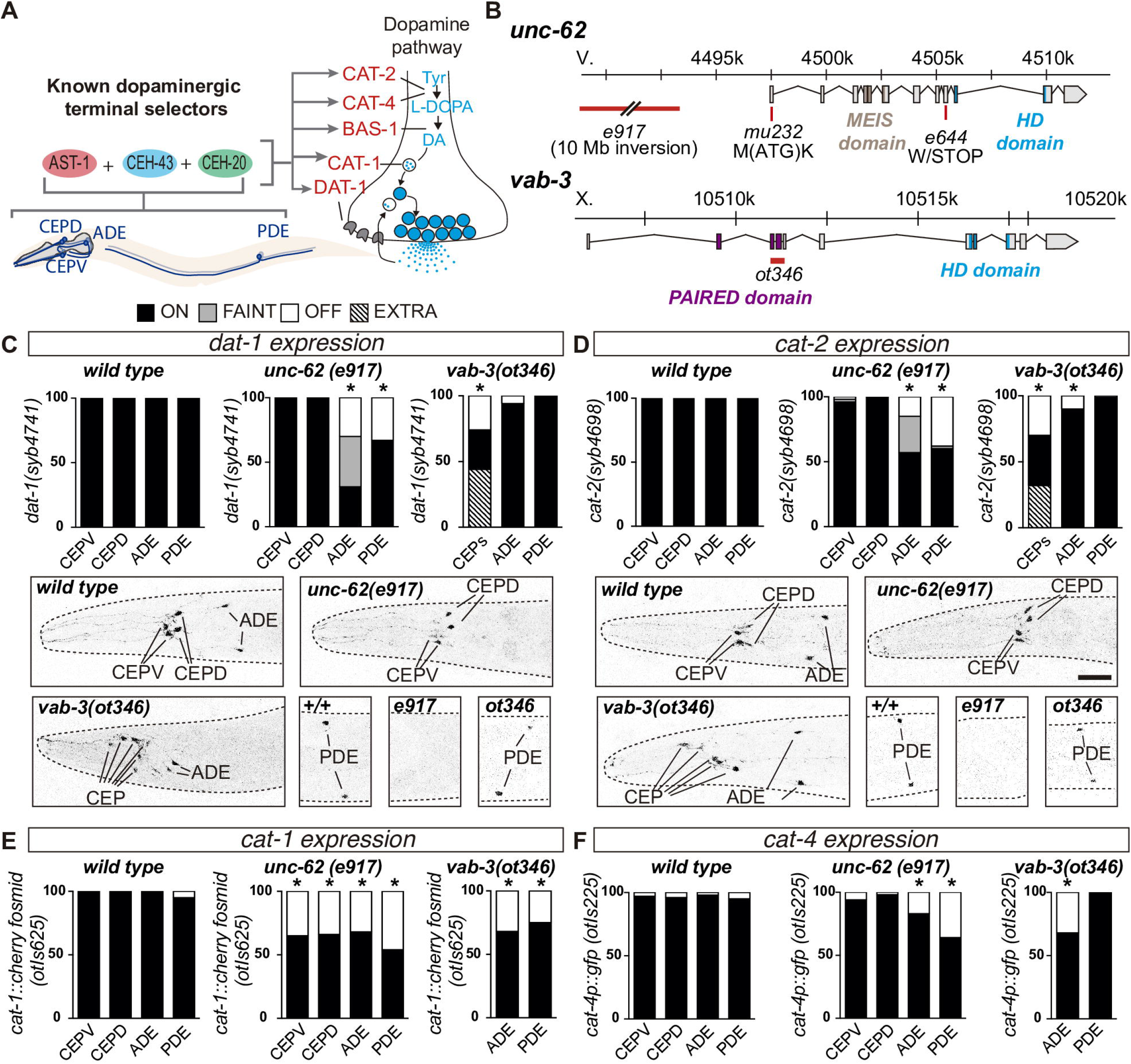
*unc-62/MEIS* HD and *vab-3*/PAIRED HD are required for correct dopamine pathway gene expression in all dopaminergic subtypes. **A**) AST-1/ETS, CEH-43/DLL HD and CEH-20/PBX HD are known terminal selectors for all four dopaminergic neuron types and directly activate expression of genes coding for the dopamine pathway components. CAT-1/VMAT1/2: vesicular monoamine transporter, CAT-2/TH: tyrosine hydroxylase, CAT-4/GCH1: GTP cyclohydrolase, BAS-1/DDC: dopamine decarboxylase, DAT-1/DAT: Dopamine transporter, DA: dopamine, Tyr: tyrosine. **B**) Schematic representation of *unc-62* and *vab-3* gene loci and alleles used in the analysis. **C-D**) Endogenous *dat-1* and *cat-2* dopamine pathway gene reporter expression analysis in *unc-62*(*e971*) and *vab-3*(*ot346*) alleles. For *cat-2* and *dat-1* analysis, disorganization of *vab-3*(*ot346*) head neurons precluded us from distinguishing CEPV from CEPD and thus are scored as a unique CEP category. Reporter expression quantification and representative micrographs of each genotype are shown. n>50 animals each condition, *: p<0.05 compared to *wild type*. Fisher’s exact test. Scale: 25 μm. **E**) *cat-1 (otIs625*) fosmid recombineered reporter (integrated multicopy array) expression analysis in *unc-62*(*e971*) and *vab-3*(*ot346*) alleles. Disorganization of *vab-3*(*ot346*) head precluded the identification of CEPs among other mCherry expressing neurons in the region and thus only ADE and PDE scoring is shown. n>50 animals each condition, *: p<0.05 compared to *wild type*. **F**) *cat-4*(*otIs225*) transcriptional reporter (integrated multicopy array) expression analysis in *unc-62*(*e971*) and *vab-3*(*ot346*) alleles. Disorganization of *vab-3*(*ot346*) head precluded the identification of CEPs among other GFP expressing neurons in the region and thus only ADE and PDE scoring is shown. n>50 animals each condition, *: p<0.05 compared to *wild type*.

### *unc-62*/MEIS-HD and *vab-3*/PAIRED-HD TFs have dual roles in dopaminergic lineage specification and terminal differentiation

*unc-62*, a MEIS HD TF, has multiple functions in development. Null alleles are embryonic lethal precluding analysis of dopaminergic differentiation defects (Van Auken et al. 2002). Three viable hypomorphic alleles *e644*, *mu232* and *e917* show expression defects of a *cat-2*/tyrosine hydroxylase reporter, the rate-limiting enzyme for dopamine synthesis (**Fig. 2B** and **Supplemental Fig. S5**). To further characterize additional dopamine pathway genes, we focused on *unc-62(e917*) allele since it showed higher penetrance of missing reporter expression. Reporter expression of endogenously tagged *cat-2/tyrosine hydroxylase* and *dat-1*/*dopamine transporter* loci in *unc-62*(*e917*) mutants is affected in ADE and PDE neurons while fosmid recombineered *cat-1/vesicular MA transporter* reporter expression defects are found for all dopaminergic subtypes (**Fig. 2C-E**). Multicopy transcriptional reporters for other dopamine pathway genes or additional effector genes not directly related to dopaminergic biosynthesis, such as the ion channel *asic-1*, are also affected in *unc-62*(*e917*) mutants, predominantly in ADE and PDE neurons (**Fig. 2F** and **Supplemental Fig. S5**). *vtIs1*(*dat-1p::gfp*) reporter expression in the ADE is unaffected in *unc-62*(*e917*) mutants revealing the presence of the cell and suggesting that the ADE lineage is unaffected in this particular allele. In contrast, *unc-62*(*e917*) shows similar loss of PDE expression for all analyzed reporters which could reflect a loss of the PDE cell due to early lineage defects. In addition, we found correlated loss of ciliated marker *ift-20* and *dat-1* reporters in the PDE, and loss of panneuronal marker *rab-3* reporter in PDE and its sister cell PVD (**Supplemental Fig. S6)**. Expression of genes coding for cilia and panneuronal components are not usually affected in terminal selector mutants (Flames and Hobert 2009; Stefanakis et al. 2015), thus PDE phenotypes in *unc-62*(*e917*) are in agreement with an early role for *unc-62* in correct PDE lineage generation. Finally, expression of two PVD markers *asic-1* reporter (also expressed in PDE) and *unc-86* TF reporter (expressed in PVD and not PDE) are also affected in *unc-62*(*n917*) mutants (**Supplemental Fig. S2**). Altogether, our results are consistent with a role for *unc-62* in CEPs and ADE terminal differentiation and with either broad expression defects in PDE and PVD cells or defects in V5 lineage generation. The potential *unc-62* role in PDE lineage formation precludes the assignment of later functions, however adult animals express *unc-62* both in ADE and PDE (Reilly et al. 2020) which could indicate a role in the terminal differentiation for both cell types.

To characterize the role of *vab-3* in dopaminergic terminal differentiation we analyzed *vab-3*(*ot346*), a deletion allele originally isolated from a forward genetic screen for dopaminergic mutants (**Fig. 2B**) (Doitsidou et al. 2008). Similar to our RNAi results, reported defects in *vab-3*(*ot346*) consists of a mixed phenotype of *vtIs1*(*dat-1p::gfp*) reporter extra and missing CEPs and, accordingly, *vab-3* was proposed to act as an early determinant of CEP lineages (Doitsidou et al. 2008). In addition to this phenotype, *vab-3*(*ot346*) expression analysis of all dopamine pathway gene reporters, including endogenously tagged *dat-1/dopamine transporter* and *cat-2/tyrosine hydroxylase* genes unravels expression defects in all dopaminergic subtypes (**Fig. 2 C-F** and **Supplemental Fig. S5)**. Of note, *dat-1* endogenous reporter expression is unaffected in ADE and PDE, which strongly suggests that the missing expression phenotype for other gene reporters in these two neuron types is not due to absence of the cells themselves (**Fig. 2C, E**). We confirmed CEP lineage defects in *vab-3(ot346*) mutants by analyzing *gcy-36* and *mod-5* reporter expression which are effector genes expressed in URX (sister cell of CEPD) and AIM (cousin of CEPV) respectively. We found that both are ectopically expressed in *vab-3*(*ot346*) animals (**Supplemental Fig. S6**).

Thus, similar to *unc-62*, *vab-3* seems to have a dual role in dopaminergic specification: it is required for proper ADE and PDE terminal differentiation and for correct CEP lineage generation. A potential role for *vab-3* in CEPs terminal differentiation could be masked by its earlier requirement in lineage determination. It has been reported that adult worms maintain *vab-3* expression in CEPs while adult ADE and PDE expression has not been reported (Reilly et al. 2020).

### PAX6, mouse ortholog of VAB-3 rescues *vab-3*(*ot346*) expression defects

Mouse orthologs for the known dopaminergic terminal selectors *ast-1*, *ceh-43* and *ceh-20* (Mouse Etv1, Dlx2 and Pbx1 respectively) are required for mouse olfactory bulb dopaminergic terminal differentiation (Brill et al., 2008; Cave et al., 2010; Flames and Hobert, 2009; Remesal et al., 2020). Thus, the dopaminergic terminal differentiation program seems to be phylogenetically conserved. Meis2 (the mouse ortholog of *unc-62*) and Pax6 (the mouse ortholog of *vab-3*) are also necessary for olfactory bulb dopaminergic specification (Agoston et al., 2014; Brill et al., 2008).

To further study functional conservation of the dopaminergic regulatory program, we performed rescue experiments for *vab-3* mutants using both the *C. elegans* gene and the mouse ortholog Pax6. Expression under the *dat-1* promoter (unaffected in ADE) of the *vab-3* isoform a mRNA, containing both the PAIRED and the HD DNA-binding protein domains, is sufficient to rescue *cat-2* reporter expression defects in ADE neuron, demonstrating a cell autonomous and terminal role for *vab-3* in dopaminergic neuron specification (**Supplemental Fig. S7**). Similarly, mouse Pax6 is able to rescue *vab-3* phenotype (**Supplemental Fig. S7**). As expected, *vab-3*(*ot346*) CEP lineage defects were not rescued, as the promoter used for rescue (*dat-1prom*) is only expressed in terminally differentiated CEPs. These data support the phylogenetic conservation of the gene regulatory network controlling dopaminergic terminal differentiation.

### UNC-62 and VAB-3 are required to maintain dopaminergic effector gene expression

To avoid early progenitor defects in *unc-62*(*e917*) and *vab-3*(*ot346*) mutants and to study the cell autonomous role of these TFs in dopaminergic neurons we performed cell specific RNAi experiments expressing double stranded RNA under a *dat-1* promoter. *unc-62*(*dat-1pRNAi*) animals lose *cat-2/tyrosine hydroxylase* endogenous expression in all dopaminergic subtypes and show reduced expression of endogenous *dat-1/dopamine transporter* (**Fig. 3A, B**). Faint expression suggests cells are still present but fail to properly maintain dopamine pathway gene expression. Similar results were found in *vab-3*(*dat-1pRNAi*) animals **(Fig. 3A, B)**. In contrast to *vab-3*(*ot346*) mutant phenotype, no ectopic reporter expression is found in *vab-3*(*dat-1pRNAi*) animals as would be expected from the specific loss of *vab-3* in the postmitotic dopaminergic neurons but not earlier in the lineage or in other postmitotic cells.

**Figure 3.**
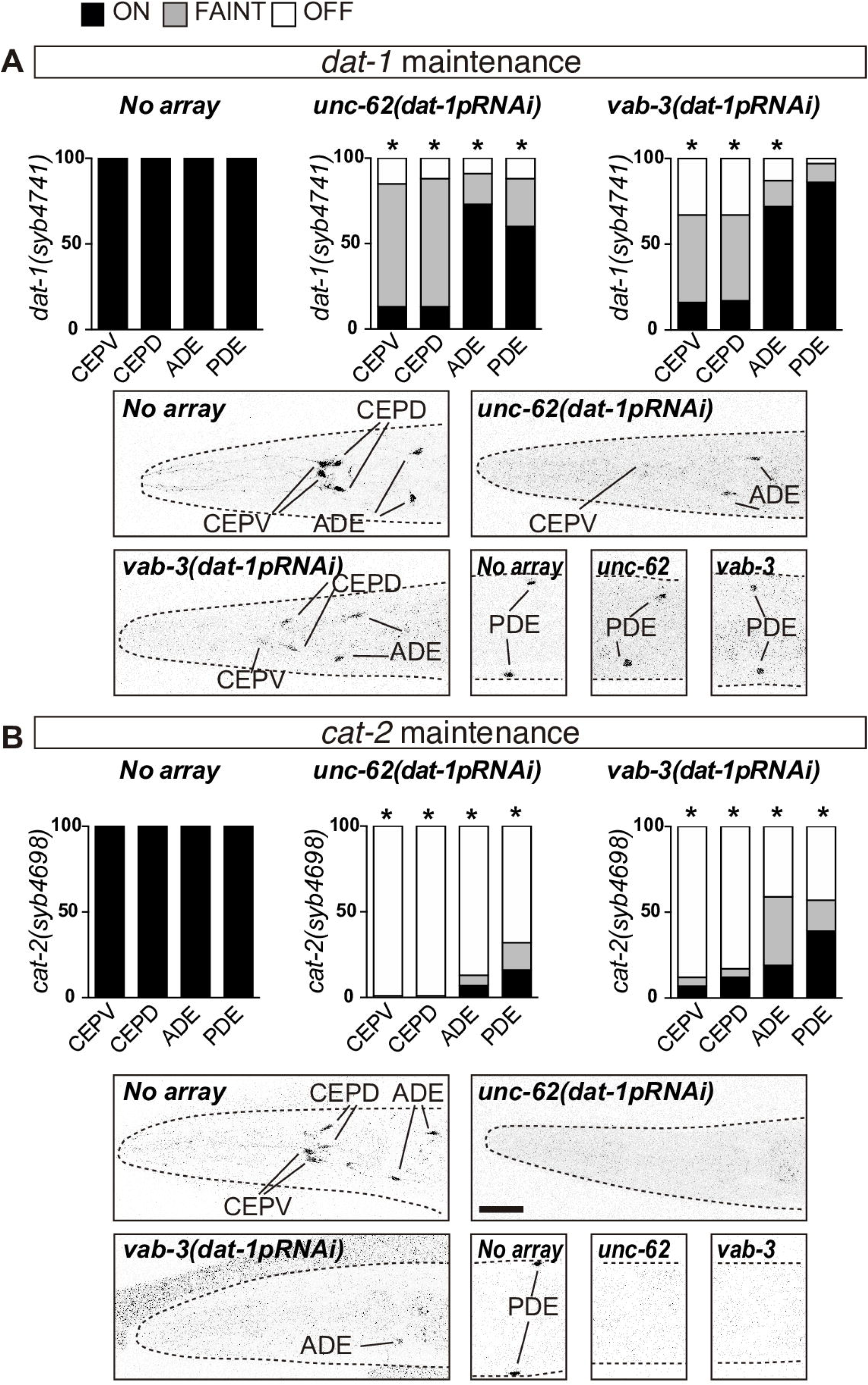
*unc-62* and *vab-3* dopaminergic specific RNAi induce defects in endogenous *dat-1* and *cat-2* gene expression maintenance. **A-B**) Endogenous *dat-1* and *cat-2* dopamine pathway gene reporter expression analysis in animals expressing *unc-62* and *vab-3* double stranded RNA under the dopaminergic specific promoter *dat-1*. Reporter expression quantification and representative micrographs of each genotype are shown. n>50 animals each condition, *: p<0.05 compared to *wild type*. Fisher’s exact test. Scale: 25 μm.

Altogether, cell type specific RNAi experiments are consistent with a terminal role for *unc-62* and *vab-3* TFs in dopaminergic terminal differentiation and effector gene expression maintenance.

### Functional binding sites for UNC-62 and VAB-3 are required for correct *cat-2/tyrosine hydroxylase* dopaminergic effector gene expression

We previously reported functional binding sites (BS) for *ast-1*, *ceh-43* and *ceh-20* terminal selectors in the *cis*-regulatory modules of the dopamine pathway genes (Flames and Hobert 2009; Doitsidou et al. 2013). To analyze whether *unc-62* and *vab-3* also directly activate dopaminergic effector gene expression, we focused on the analysis of *cis* features regulating *cat-2/tyrosine hydroxylase* that is exclusively expressed in the dopaminergic neurons and severely affected by *unc-62* and *vab-3* loss.

*cat-2* minimal *cis*-regulatory module (*cat-2p21*), in addition to the previously described ETS (*ast-1*), PBX (*ceh-20*) and HD (*ceh-43*) functional BS (Flames and Hobert 2009; Doitsidou et al. 2013), also contains predicted MEIS and PAIRED BS (**Fig. 4A**). Extrachromosomal reporter constructs of *cat-2* minimal promoter with point mutations in PAIRED and MEIS BS show strong *gfp* expression defects in all dopaminergic neurons except CEPV (**Fig. 4A**).

**Figure 4.**
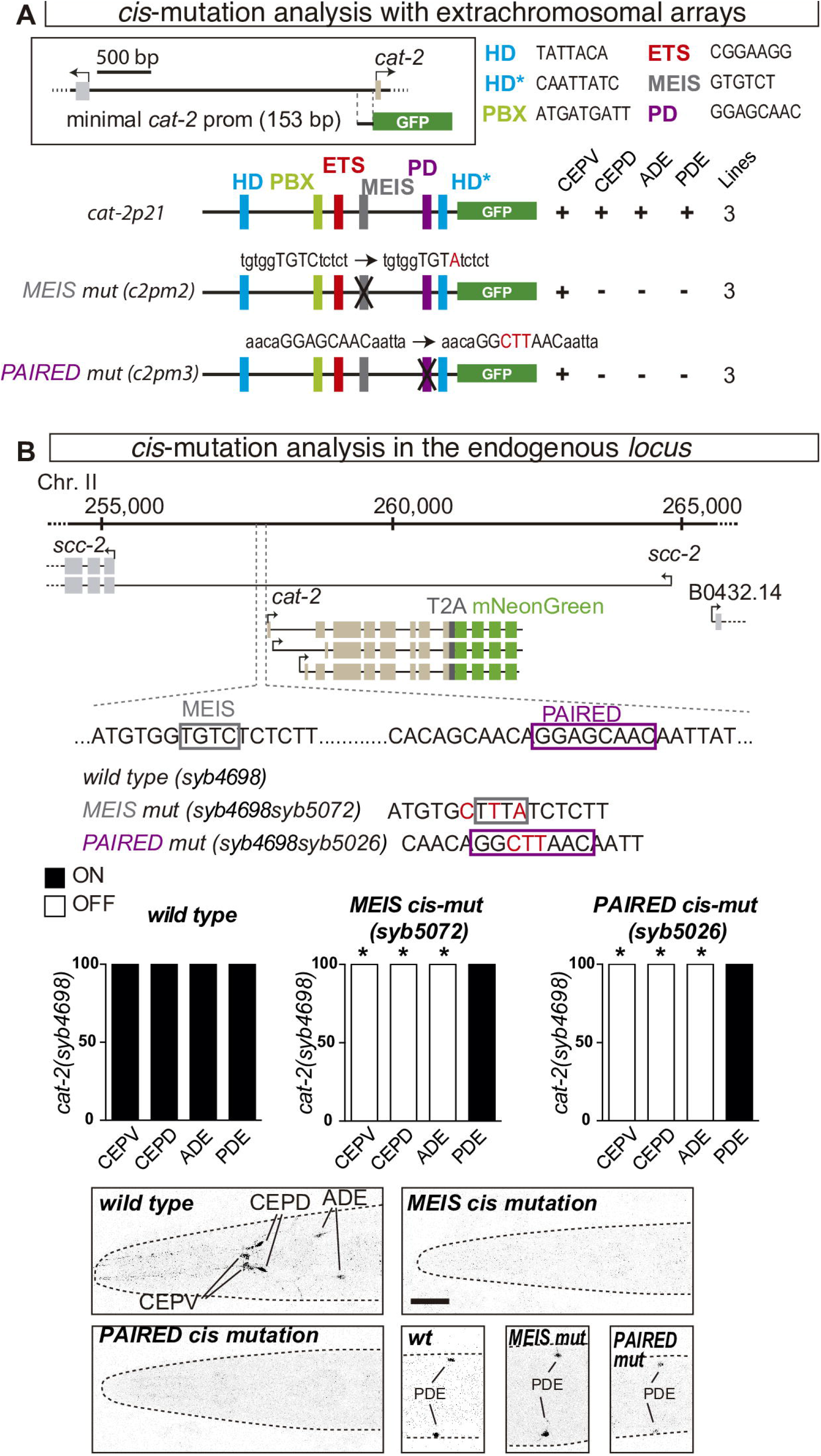
*cis*-regulatory analysis of *cat-2/tyrosine hydroxylase* effector gene reveals functional binding sites for UNC-62 and VAB-3. **A**) Multicopy extrachromosomal reporter analysis of *cat-2* minimal dopaminergic *cis*-regulatory module (*cat-2p21*). In addition to published functional AST-1/ETS, CEH-20/PBX HD, CEH-43/DLL HD, binding sites (Doitsidou et al. 2013; Flames and Hobert 2009), UNC-62/MEIS and VAB-3/PAIRED binding sites are also required for correct GFP reporter expression in different dopaminergic neuron types. HD* represents a PAIRED-type HD consensus (HTAATTR). Black crosses represent point mutations to disrupt the corresponding TFBS, the nature of the mutation is indicated in red. +: > 70% of mean *wild type* construct values; -: values are less than 10% expression in at least two of the three analyzed lines. n > 30 animals per line. **B**) Analysis of the effect of MEIS and PAIRED TFBS *cis* mutations in the expression of endogenous *cat-2* gene. Schema of used strains with the corresponding mutations, quantification of reporter gene expression defects and representative micrographs are shown. n>50 animals each condition, *: p<0.05 compared to *wild type*. Scale: 25 μm

VAB-3 protein contains two DNA-binding domains, PAIRED and HD, each fulfilling specific functions (Brandt et al. 2019). We found that one of the two already identified functional HD BS (Doitsidou et al. 2013) matches a PAIRED-type HD consensus motif (HTAATTR, labeled as HD* in **Fig. 4A**) suggesting it could be recognized by VAB-3 and/or CEH-43.

Next, we studied the effect of MEIS and PAIRED BS mutations in the context of the *cat-2* endogenous locus (**Fig. 4B**). Both mutations abolish *cat-2* gene expression in CEPD and ADE, similar to effects found with the extrachromosomal arrays. However, CEPV *cat-2* expression is also affected in these animals, while PDE expression is unaffected. Discrepancies between *cis* regulatory mutations in extrachromosomal and the endogenous locus suggest chromatin context and epigenetic modifications might modulate the effect of *cis* mutations in some dopaminergic subtypes but not in others.

### The dopaminergic regulatory signature is preferentially associated to dopaminergic neuron effector genes

Our results show that at least five TFs seem to be required for correct *C. elegans* dopaminergic terminal specification. These findings coincide with a seemingly complex regulatory logic found in HSN serotonergic neurons (Lloret-Fernández et al. 2018). We previously described that TF binding site (TFBS) clusters for the HSN TF collective are preferentially found in regulatory regions near HSN expressed genes and can be used to *de novo* identify enhancers active in the HSN neuron (Lloret-Fernández et al. 2018). Thus, a function for this complex terminal differentiation programs might be to provide enhancer selectivity.

To test this hypothesis, we asked if the dopaminergic regulatory program imposes a regulatory signature in dopaminergic-expressed genes. Published single-cell RNA-seq data (Cao et al. 2017) was used to identify additional genes differentially expressed in dopaminergic neurons (**Supplemental Fig. S8**). We found 86 genes whose expression is enriched in dopaminergic neurons compared to other clusters of ciliated sensory neurons. As expected, this gene list includes all dopamine pathway genes and other known dopaminergic effector genes, but not pancilia expressed genes (**Supplemental Table S7**). In analogy to our previous analysis of the HSN regulatory genome (Lloret-Fernández et al. 2018), we analyzed the upstream and intronic sequences of these genes. For comparison purposes, we built ten thousand sets of 86 random genes with similar upstream and intronic length distribution to dopaminergic expressed genes.

First, we focused our analysis only on the three already published dopaminergic terminal selectors (AST-1/ETS, CEH-43/ HD and CEH-20/PBX HD). This simple regulatory signature (presence of ETS+HD+PBX binding sites clustered in 700 bp or smaller regions) lacks specificity as all genes, either dopaminergic expressed genes or random sets, contain DNA windows with matches for all three TFs (100% of dopaminergic expressed genes compared to 100% in random sets). Reducing DNA-window search length from 700 bp to 300 bp or 150 bp did not increase specificity of dopaminergic signature in dopaminergic expressed genes (**Supplemental Table S7**). Next, we expanded our analysis to the search of a regulatory signature including UNC-62/MEIS and VAB-3/PAIRED and HD* predicted binding sites (ETS+HD+HD*+PBX+MEIS+PAIRED binding sites clustered in 700bp or smaller region) (**Fig. 5A**). We found that 88% of dopaminergic expressed genes contain at least one associated dopaminergic regulatory signature window which is a percentage only slightly higher than the mean percentage found for the 10,000 random sets (84%) (p=0.4027, Chi square test) (**Fig. 5B**). Next, we built similar sets of differentially expressed genes for five randomly picked non-dopaminergic neuron categories (RIA, ASE, Touch Receptor neurons, GABAergic neurons and ALN/PLN/SDQ cluster) (**Supplemental Fig. S8** and **Supplemental Table S7**). We found that the percentage of expressed genes containing the dopaminergic signature is smaller in non-dopaminergic neurons compared to dopaminergic neurons (**Fig. 5B**). This difference is statistically significant for gene sets expressed in ASE and RIA neuron but not compared to other neuron categories (**Fig. 5B**).

**Figure 5.**
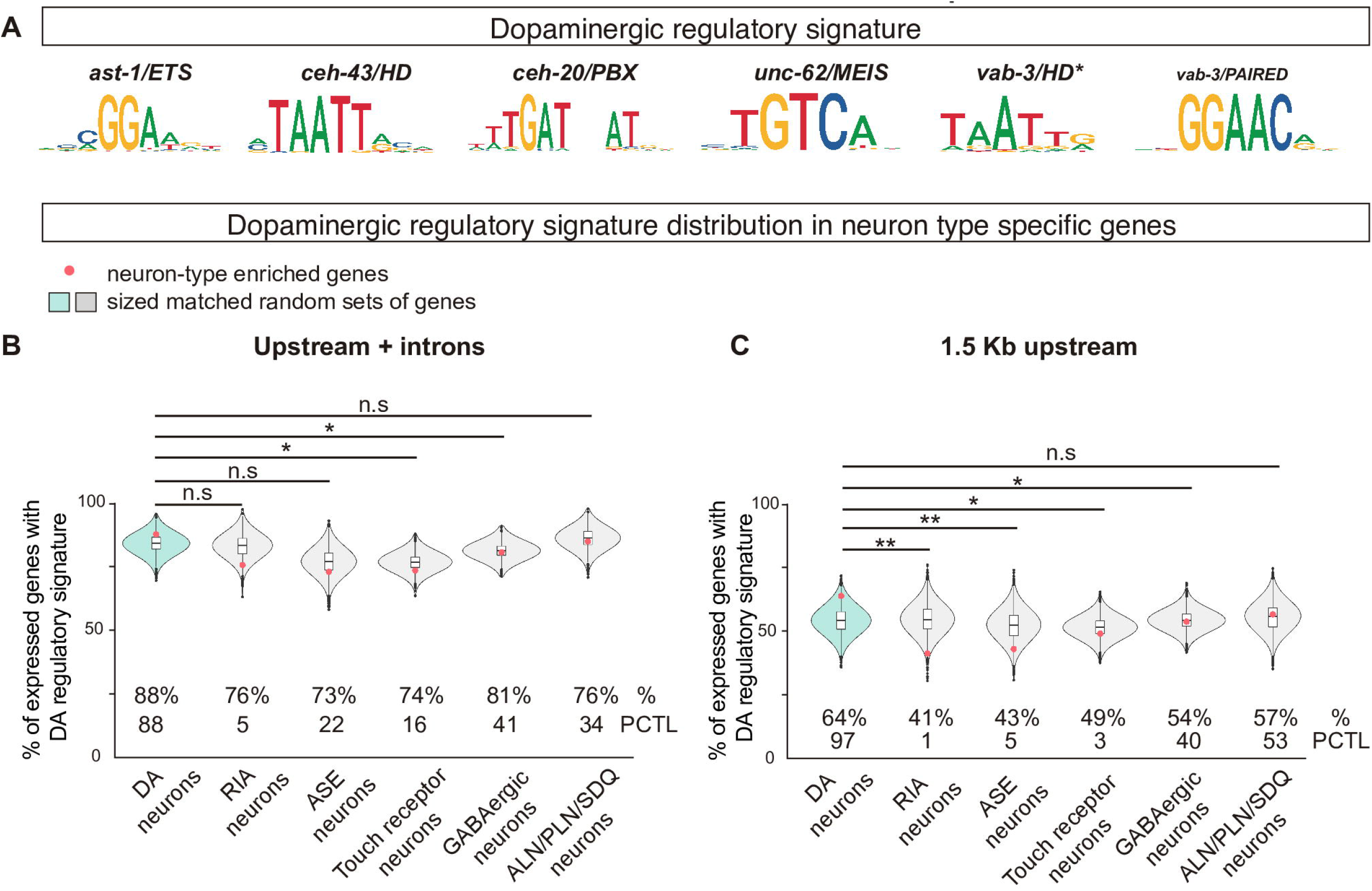
The Dopaminergic Regulatory Signature is preferentially associated to dopaminergic expressed genes. **A**) Position weight matrix logos assigned to each member of the dopaminergic terminal selectors. The dopaminergic regulatory signature is defined by the presence of at least one match for each of the six PWM in less than 700 bp DNA window. **B**) Dopaminergic regulatory signature is slightly more prevalent in the upstream and intronic sequences of the set of 86 genes with enriched expression in dopaminergic neurons (red dot in blue violin plot) compared to five additional gene sets with enriched expression in non-dopaminergic neurons (RIA, ASE, Touch receptor neurons, GABAergic neurons and ALN/PLN/SDQ). However, dopaminergic signature presence in dopaminergic enriched genes is not higher than the mean of 10,000 sets of random comparable genes (red dot location inside the blue violin plot). %: percentage of genes with assigned dopaminergic regulatory signature. PCTL: percentile of the real value (red dot) in the 10,000 random set value distribution. Brunner-Munzel test. *: p<0.05, **: p<0.01. **C**) Dopaminergic regulatory signature in proximal regions (1.5 kb upstream ATG) is more highly enriched in dopaminergic expressed genes compared to other non-dopaminergic expressed genes, and dopaminergic signature presence in dopaminergic enriched genes is higher than the mean of 10,000 sets of random comparable genes (red dot location inside the blue violin plot) suggesting proximal regulation has a major role in dopaminergic terminal differentiation. See (B) for abbreviations and statistics.

Next, we restricted dopaminergic regulatory signature search to regions proximal to the translation start site (1.5 kb upstream of the ATG). 64% of dopaminergic-enriched genes contain the dopaminergic regulatory signature in this proximal region, a higher percentage compared to the mean percentage of all random sets of genes (54%, p=0.088, Chi square test) (**Fig. 5C**). Moreover, the percentage of genes with proximal dopaminergic signature is significantly smaller in all non-dopaminergic neuronal categories except for ALN/PN/SDQ cluster. The reason why ALN/PLN/SDQ expressed genes show a higher association to dopaminergic regulatory signature compared to other non-dopaminergic neurons is uncertain. Terminal selectors for ALN/PLN/SDQ neurons are yet unknown, however, our previous work determined that SDQ neurons co-express AST-1, CEH-43 and CEH-20 (Doitsidou et al. 2013) and single cell data shows *unc-62* expression in this neuron as well (Taylor et al. 2021). Thus a similar combination of TFs controlling terminal differentiation of SDQ could explain the higher presence of the dopaminergic regulatory signature. Finally, in contrast to the random set of genes for the dopaminergic expressed genes, none of the non-dopaminergic neuronal types showed a significant enrichment of the dopaminergic signature in respect to their corresponding background of ten thousand random sets of comparable genes (**Fig. 5C**). Thus, dopaminergic regulatory signature seems to be enriched in the proximal regions of dopaminergic effector genes. Four out of five dopamine pathway genes contain at least one associated dopaminergic regulatory signature window closer than 1.5 kb from their ATGs. Experimentally isolated *cis*-regulatory modules (ranging from 143 bp to 521 bp in size) (Flames and Hobert 2009) overlap with predicted dopaminergic regulatory signature windows and contain at least one match for each of the six types of TFBS (**Supplemental Fig. S8**), the exception being *bas-1*/dopamine decarboxylase *cis*-regulatory module that lacks HD and MEIS predicted binding sites (**Supplemental Fig. S8**). TFs collectives can be recruited to target enhancers even with partial complements of their respective TFBS (Spitz and Furlong 2012). Thus we performed similar bioinformatic analysis for dopaminergic signature enrichment with partial complements of the signature (at least 5 or at least 4 TFBS classes). Flexible dopaminergic regulatory signature is still enriched in dopamine expressed genes, albeit at lower levels of significance than the complete regulatory signature, compared to random controls and is more frequent than in genes expressed in other neurons, particularly in regions located 1.5 kb or closer to ATG, (**Supplemental Fig. S9**).

Finally, we built reporter constructs to test dopaminergic regulatory signature windows from 14 non-coding regions associated to the dopamine enriched gene list. Six out of the fourteen tested constructs (43%) drive expression in the dopaminergic neurons (**Fig. 6A** and **Supplemental Table S8**) while the rest of the constructs drive reporter expression in other neurons or tissues but not in dopaminergic neurons. Dopaminergic active enhancers are located closer to ATG than reporters not expressed in dopaminergic neurons, in agreement to our bioinformatics enrichment analysis **Fig. 6B**).

**Figure 6.**
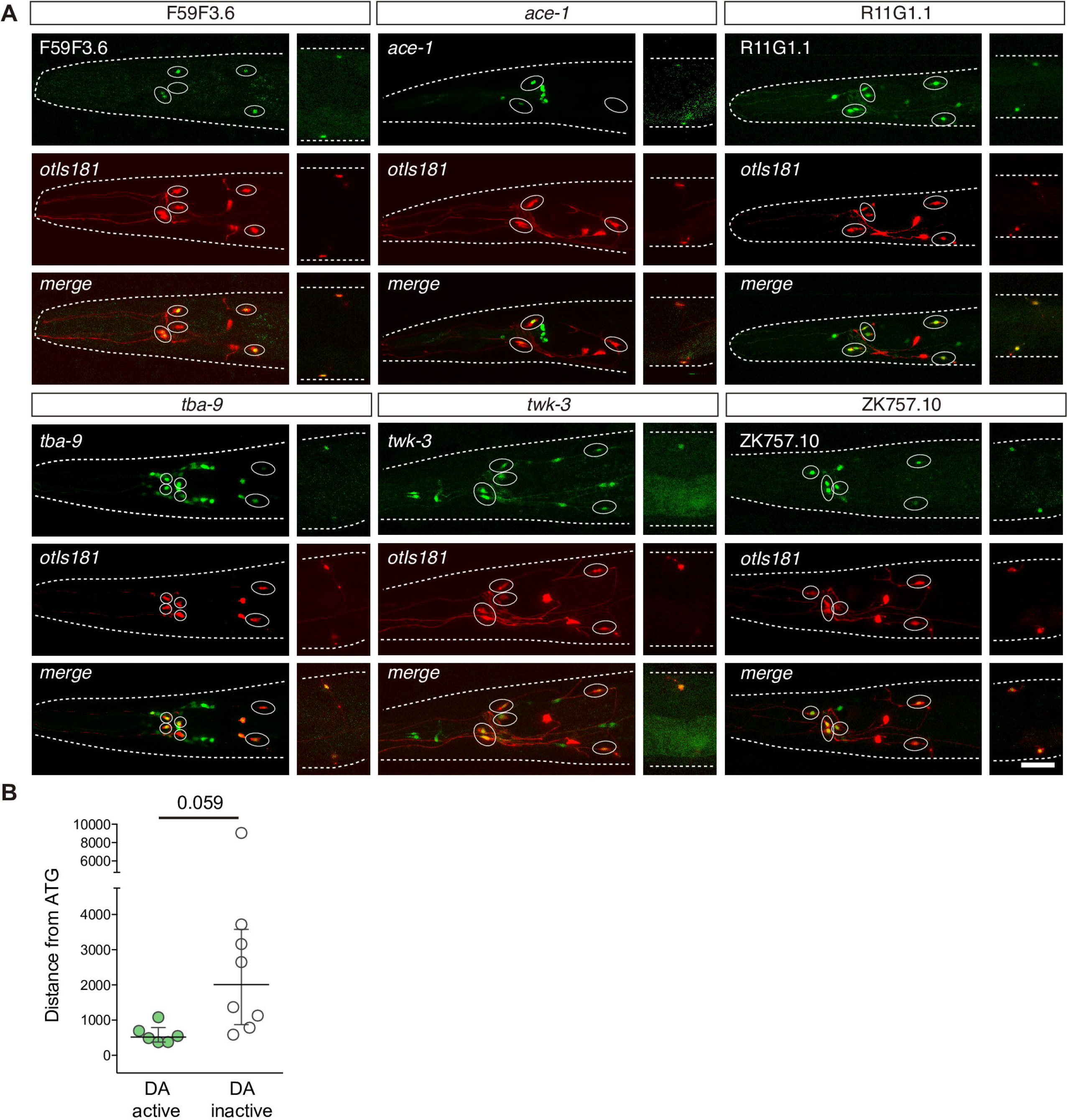
Experimental validation of dopaminergic regulatory signature. **A**) Representative micrographs showing GFP expression driven by six different genomic regions overlapping predicted dopaminergic regulatory signature windows. Dopaminergic expression was assessed by co-localization with *otIs181* reporter (*dat-1p::mcherry*; *ttx-3p::mcherry*) shown in red. Except for F59F3.6, which is exclusively expressed in dopaminergic neurons, GFP expression was also found in additional unidentified and reporter-specific neurons in the head. See **Supplemental Table S8** for quantification of two independent reporter lines for each construct. Scale: 25 μm **B**) Distance from the central location of the construct to ATG. Constructs driving GFP expression in dopaminergic neurons tend to locate at closer distances to the ATG than those not active in dopaminergic neurons. Two-tailed *t*-test.

These data concur with our previous results obtained for the HSN regulatory signature (37% success rate) (Lloret-Fernández et al. 2018) and reveal that the presence of a dopaminergic regulatory signature, particularly when located in proximal regions to the ATG, can be used to identify dopaminergic active enhancers although signature itself, as it is presently characterized, is not sufficient to induce dopaminergic expression.

### Dopaminergic regulatory signature distribution in other genes expressed in the dopaminergic neurons

Next, we aimed to analyze the presence of a dopaminergic regulatory signature in additional genes active in dopaminergic neurons. Different hierarchies of gene expression co-exist in any given cell, mechanosensory dopaminergic neurons co-express at least four types of genes: 1) Dopaminergic effector genes such as dopaminergic pathway genes, neuropeptides, neurotransmitter receptors, etc., that are preferentially (but not exclusively) expressed by this neuron type; 2) Genes coding for structural components of cilia which are expressed by all sixty ciliated sensory neurons in *C. elegans*; 3) Panneuronal genes, such as components of the synaptic machinery or cytoskeleton, expressed by all 302 neurons in *C. elegans* hermaphrodite and 4) Ubiquitous genes expressed by all cells of the organism, such as ribosomal or heat shock proteins. We found that the percentage of genes with dopaminergic signature is highest for dopaminergic enriched effector genes compared to other sets of genes both when considering complete non-coding sequences or only the proximal regions (**Supplemental Fig. S10**). Of note, albeit lower than dopaminergic effector gene set, panneuronal genes and cilia related components also showed increased presence of dopaminergic regulatory signature compared to ubiquitous or not-neuronal expressed genes (**Supplemental Fig. S10**).

## DISCUSSION

### Neuron-type specification is mainly controlled by five transcription factor families

Focusing mainly on the MA superclass of neurons we provide a comprehensive view of the TFs required in the generation of different neuronal types. Identified TFs could act at very different steps and be involved in distinct processes such as cell division, lineage commitment to neuronal fate, lineage specification, terminal differentiation or cell survival. We did not find any global regulator of MA fate, which is likely due to the molecular and functional diversity found in this superclass of neurons. However, it is also possible that genetic redundancy, early larval lethality or false negatives of our experimental approach could preclude us from identifying pan-MA regulators.

Each neuron type is regulated by different sets of TFs, however, they consistently belong to the five most prevalent *C. elegans* TF families: HD, bHLH, ZF, bZIP and phylogenetically conserved members of the NHR family. Analysis of genetic mutants displaying diverse neuronal phenotypes in *C. elegans*, mouse and *Drosophila* reveals the same TF family distribution, which validate the results obtained from the RNAi screen and expands these findings to other neuronal types and organisms.

We find HD TFs are enriched in *C. elegans* and mouse TF neuronal mutants compared to genomic distribution. Despite the prevalent role of HD TFs as terminal selectors (Hobert 2016), these TFs also play earlier roles in progenitor specification, such as the pleiotropic actions of UNC-86 POU HD TF (Leyva-Díaz et al. 2020; Finney and Ruvkun 1990). Our data suggest *vab-3*/PAIRED HD and *unc-62*/MEIS HD TFs could also play dual roles both in progenitors and postmitotic dopaminergic neurons. It has been recently proposed that ancestral homeobox genes could be responsible for the regulation of the ancestral neuron types, and that this functional linkage has been maintained and diversified throughout evolution (Durham et al. 2021).

In addition to HD TFs, several NHRs, bZIP and ZF TFs are known to regulate neuron-type identities in mammals, such as Couptf1, Nurr1, Tlx and Nr2e3 NHR TFs (Bovetti et al. 2013; Haider et al. 2000; Zetterström et al. 1997; Roy et al. 2004), Nrl, cMaf and Mafb bZIP TFs (Wende et al. 2012; Blanchi et al. 2003; Mears et al. 2001) or Myt1l, Gli1, Sp8, Ctip2, Fezf1/2 ZF TFs (Mall et al. 2017; Hynes et al. 1997; Waclaw et al. 2006; Arlotta et al. 2005; Shimizu et al. 2010). In contrast to known proneuronal actions for bHLH and main role in establishing specific indentities for HD, described for different neuron types and species, a possible general role for these additional TF families has been poorly studied in any model organism. Functional characterization of the NHR, bZIP and ZF TF candidates retrieved from our RNAi screen will help better understand the role of these TF families in the establishment of specific neuronal identities.

### Global dopaminergic regulatory logic versus dopaminergic subtype identities

The four dopaminergic neuron subtypes arise from different progenitors and display distinct morphologies, however they all converge to fulfill the same mechanosensory functions and to signal through the dopamine neurotransmitter. Accordingly, many of the genes expressed by dopaminergic mature neurons are shared among subtypes and for their activation they converge into the same regulatory program of terminal selectors. Our data, together with previous work (Flames and Hobert 2009; Doitsidou et al. 2013), suggest that the dopaminergic terminal selector collective acts through the dopaminergic regulatory signature to activate expression of general dopaminergic effector genes. TF mutant phenotypes, *cis* mutations or TF expression are often more pronounce in specific subtypes of dopaminergic neurons (**Supplemental Table S9**). For example, expression defects for some effector genes in *vab-3* mutant are higher in CEPs than in ADE and PDE and conversely, *ceh-20* and *unc-62* mutations affect more broadly (although not exclusively) PDE gene expression [this work and (Doitsidou et al. 2013)]. These data suggest that global actions of the dopaminergic terminal selector collective can be modulated in a subtype-specific manner, maybe by the action of additional subtype specific TFs.

Moreover, some effector genes are expressed only in specific dopaminergic neuron subtypes (such as neuropeptide *flp-33* expression in ADE and PDE neurons). Further investigation is required to assess if the dopaminergic terminal selector collective is also involved in dopaminergic subtype gene expression and/or if additional TFs act as subtype specific activators or repressor in dopaminergic neurons, as has been previously determined for motoneuron diversification in *C. elegans* (Kerk et al. 2017; Kratsios et al. 2017). Finally, in addition to dopaminergic effector genes, all dopaminergic neurons share the expression of other genes that are more broadly expressed, such as cilia components (ciliome) expressed by all sensory ciliated neurons, panneuronal genes shared by all neurons or ubiquitous genes. It is still unclear to what extent terminal selectors participate in the direct activation of these genes. Enhancers of panneuronal genes are very redundant and are to some extent regulated by terminal selectors (Stefanakis et al. 2015), but additional TFs might regulate their expression. Genes coding for the ciliome are directly regulated by the RFX TF DAF-19 (Swoboda et al. 2000), however DAF-19 is also expressed in non-ciliated neurons suggesting also additional TFs might participate in ciliome regulation. Our data shows that the dopaminergic regulatory signature is most abundant in dopaminergic effector genes, however panneuronal and panncilia genes show a higher percentage of dopaminergic signature than ubiquitous or non-neuronal genes suggesting dopaminergic terminal selectors could also have a direct role in their regulation.

### Complexity of terminal differentiation programs provide enhancer selectivity

The sequence determinants that differentiate active regulatory regions from other non-coding regions of the genome are currently largely unknown. To date, one of the best sequence predictors of active enhancers is the number of putative TF binding sites for different TFs found in a region (Kheradpour et al. 2013; Tewhey et al. 2016; Grossman et al. 2017). Here we have characterized the role of five different TFs that work together to direct dopaminergic terminal differentiation. Although the majority of these TFs belong to the HD family, three of the four HD members of the dopaminergic terminal selector collective contain additional DNA-binding domains that increase the variety of recognized consensus binding sites: the PBC domain of the PBX TF CEH-20, the MEIS domain of UNC-62 and the PAIRED domain in VAB-3. We hypothesize that this TFBS complexity is important to achieve enhancer selectivity. Our results show that binding site clusters of the dopaminergic TF collective are preferentially located in putative regulatory regions of dopaminergic-expressed genes compared to other genes, particularly in regions near the ATG.

The presence of the signature can be used to identify enhancers active in dopaminergic neurons. However, both presence of the dopaminergic signature in genes not expressed in dopaminergic neurons and tested dopaminergic regulatory windows from dopaminergic-expressed genes that are not active in dopaminergic neurons demonstrate that the signature, as it is currently characterized, is not sufficient to induce dopaminergic expression. More restrictive PWMs for the dopaminergic TF collective, additional TFBS, gene repression mechanisms, or chromatin accessibility is likely to further regulate dopaminergic regulatory signature specificity. It is also possible that specific syntactic rules (TFBS order, distance and disposition) discriminate functional from non-functional dopaminergic regulatory signature windows. Future experiments should be aimed to increase our understanding on the regulatory rules that define dopaminergic-effector-gene active enhancers.

## METHODS

### *C. elegans* strains and genetics

*C. elegans* culture and genetics were performed as described (Brenner 1974). Strains used in this study are listed in **Supplemental Table S10**.

### Generation of the transcription factor RNAi library

To generate a complete TF RNAi library we used the *C. elegans* TFs list from (Narasimhan et al. 2015). See **Supplemental Methods** and for a description of all used sources.

### RNAi feeding experiments and cell type–specific RNAi

RNAi feeding experiments were performed following standard protocols (Kamath et al. 2001). For cell type–specific RNAi experiments, PCR fusion of *dat-1* promoter to sense or antisense *vab-3* and *unc-62* cDNA sequences were generated and injected as previously described (Esposito et al. 2007). See **Supplemental Methods** and **Supplemental Table S11** for further details on feeding protocol, primers and microinjection conditions.

### Mutant strains and genotyping

Strains used in this study are listed in **Supplemental Table S10**. Deletion alleles were genotyped by PCR. Point mutations were genotyped by sequencing (**Supplemental Table S11**). Alleles *vab-3*(*ot346*), *unc-62*(*e644*) *and lag-1*(*q385*) were determined by visual mutant phenotype. The mutation *unc-62*(*mu232*) was followed through its link with the fluorescent reporter *muIs35*.

### Generation of *C. elegans* alleles and transgenic lines

Knock-in strains PHX4698 *cat-2*(*syb4698*), PHX4741 and *dat-1*(*syb4741*) as well as the point-mutated strains PHX5072 *cat-2*(*syb4698 syb5072*) and PHX5026 *cat-2*(*syb4698 syb5026*) were generated by SunyBiotech’s CRISPR services. Gene constructs for *cis*-regulatory analysis with extrachromosomal arrays were generated by cloning into pPD95.75 vector. Directed mutagenesis was performed with Quikchange II XL site-directed mutagenesis kit (Stratagene). For rescue and cell type-specific RNAi experiments, commercially available cDNA of the TF candidates was obtained from Dharmacon Inc and Invitrogen. Reporters for the analysis of the dopaminergic regulatory signature were created by fusion PCR (Hobert 2002). See **Supplemental Methods** for more details.

### Scoring and statistics

Scoring was performed over anesthetized animals (0.5 M of sodium azide (Sigma, #26628-22-8) on 4% agarose pads) using a 40X objective in a Zeiss Axioplan 2 microscope. All micrographs showed in this paper were obtained with a TCS-SP8 Leica Microsystems confocal microscope using a 63X objective. See **Supplemental Methods** for detailed information on samples’ sizes and scoring criteria and conditions and **Supplemental Table S8** for raw scoring data and statistics.

### Bioinformatics

#### TF family distribution in different species

Data mining of different databases in *Caenorhabditis elegans, Drosophila melanogaster* and *Mus musculus* was used to retrieve lists of transcription factors (TFs) of each species with associated neuronal phenotypes, (see **Supplemental Methods**).

#### Dopaminergic signature analysis

scRNA-seq data from (Cao et al. 2017) and (Packer et al. 2019) was used to build comprehensive gene lists for different neuronal and non-neuronal categories. For dopaminergic regulatory signature analysis, we matched different PWM against putative regulatory regions in the genome using a sliding-window approach, following methodology described in (Lloret-Fernández et al. 2018). The same analysis was conducted restricting the search space up to 1.5 kb upstream the ATG of the gene. We then assessed the specificity of the dopaminergic regulatory signature in different neuron-type enriched gene lists. These analyses are further detailed in **Supplemental Methods**.

## Acknowledgments

We thank CGC (P40 OD010440) for providing strains. Elia García and Francisco Anguix for technical help. Ana Pilar Gómez-Escribano and Carla Lloret-Fernández for helping in the generation of the RNAi library. Adrián Tarazona for helping in the bioinformatics and statistical analysis. Bioinformatics and Biostatistics Unit from Principe Felipe Research Center (CIPF) for providing access to the cluster, co-funded by European Regional Development Funds (FEDER). Dr Peter Askjaer and Dr. Marta Artal for sharing reagents. Dr Guillermo Ayala for advice on statistical analysis. Dr Oscar Marín, Dr Beatriz Rico and Dr Luisa Cochella labs for scientific discussion. Dr Oscar Marín, Dr Luisa Cochella, Dr Inés Carrera, Dr Roger Pocock, Dr Arantza Barrios and Dr Oliver Hobert for comments on the manuscript. Funding sources: ERC-StG-2011-281920; ERC-Co-2020-101002203; SAF2017-84790-R; PID2020-115635RB-I00; PROMETEO/2018/055; RED2018-102553-T and ACIF/2019/079.

## REFERENCES

Agoston Z, Heine P, Brill MS, Grebbin BM, Hau AC, Kallenborn-Gerhardt W, Schramm J, Götz M, Schulte D. 2014. Meis2 is a Pax6 co-factor in neurogenesis and dopaminergic periglomerular fate specification in the adult olfactory bulb. Dev.

Andrews MG, Kong J, Novitch BG, Butler SJ. 2019. New perspectives on the mechanisms establishing the dorsal-ventral axis of the spinal cord. In Current Topics in Developmental Biology.

Angerer LM, Yaguchi S, Angerer RC, Burke RD. 2011. The evolution of nervous system patterning: Insights from sea urchin development. Development.

Arlotta P, Molyneaux BJ, Chen J, Inoue J, Kominami R, MacKlis JD. 2005. Neuronal subtype-specific genes that control corticospinal motor neuron development in vivo. Neuron 45: 207–221.

Bertrand N, Castro DS, Guillemot F. 2002. Proneural genes and the specification of neural cell types. Nat Rev Neurosci.

Blanchi B, Kelly LM, Viemari JC, Lafon I, Burnet H, Bévengut M, Tillmanns S, Daniel L, Graf T, Hilaire G, et al. 2003. MafB deficiency causes defective respiratory rhythmogenesis and fatal central apnea at birth. Nat Neurosci 6: 1091–1099.

Bodofsky S, Koitz F, Wightman B. 2017. Conserved and Exapted Functions of Nuclear Receptors in Animal Development. Nucl Recept Res 4: 1–33.

Borello U, Pierani A. 2010. Patterning the cerebral cortex: Traveling with morphogens. Curr Opin Genet Dev.

Bovetti S, Bonzano S, Garzotto D, Giannelli SG, Iannielli A, Armentano M, Studer M, De Marchis S. 2013. COUP-TFI controls activity-dependent tyrosine hydroxylase expression in adult dopaminergic olfactory bulb interneurons. Dev.

Brandt JP, Rossillo M, Du Z, Ichikawa D, Barnes K, Chen A, Noyes M, Bao Z, Ringstad N. 2019. Lineage context switches the function of a C. elegans Pax6 homolog in determining a neuronal fate. Dev.

Brenner S. 1974. The genetics of Caenorhabditis elegans. Genetics.

Brill MS, Snapyan M, Wohlfrom H, Ninkovic J, Jawerka M, Mastick GS, Ashery-Padan R, Saghatelyan A, Berninger B, Götz M. 2008. A Dlx2- and Pax6-dependent transcriptional code for periglomerular neuron specification in the adult olfactory bulb. J Neurosci.

Briscoe J, Pierani A, Jessell TM, Ericson J. 2000. A homeodomain protein code specifies progenitor cell identity and neuronal fate in the ventral neural tube. Cell.

Cao J, Packer JS, Ramani V, Cusanovich DA, Huynh C, Daza R, Qiu X, Lee C, Furlan SN, Steemers FJ, et al. 2017. Comprehensive single-cell transcriptional profiling of a multicellular organism. Science (80-) 357: 661–667. https://www.science.org/doi/10.1126/science.aam8940.

Cave JW, Akiba Y, Banerjee K, Bhosle S, Berlin R, Baker H. 2010. Differential regulation of dopaminergic gene expression by Er81. J Neurosci.

Doitsidou M, Flames N, Lee AC, Boyanov A, Hobert O. 2008. Automated screening for mutants affecting dopaminergic-neuron specification in C. elegans. Nat Methods.

Doitsidou M, Flames N, Topalidou I, Abe N, Felton T, Remesal L, Popovitchenko T, Mann R, Chalfie M, Hobert O. 2013. A combinatorial regulatory signature controls terminal differentiation of the dopaminergic nervous system in C. elegans. Genes Dev 27: 1391–1405. http://genesdev.cshlp.org/cgi/doi/10.1101/gad.217224.113.

Durham TJ, Daza RM, Gevirtzman L, Cusanovich DA, Bolonduro O, Noble WS, Shendure J, Waterston RH. 2021. Comprehensive characterization of tissue-specific chromatin accessibility in L2 Caenorhabditis elegans nematodes. Genome Res 31: 1952–1969. http://dx.doi.org/10.1038/s41583-021-00497-x.

Esposito G, Di Schiavi E, Bergamasco C, Bazzicalupo P. 2007. Efficient and cell specific knock-down of gene function in targeted C. elegans neurons. Gene 395: 170–176.

Finney M, Ruvkun G. 1990. The unc-86 gene product couples cell lineage and cell identity in C. elegans. Cell 63: 895–905. https://linkinghub.elsevier.com/retrieve/pii/009286749090493X.

Flames N, Hobert O. 2009. Gene regulatory logic of dopamine neuron differentiation. Nature.

Flames N, Hobert O. 2011. Transcriptional Control of the Terminal Fate of Monoaminergic Neurons. Annu Rev Neurosci.

Grossman SR, Zhang X, Wang L, Engreitz J, Melnikov A, Rogov P, Tewhey R, Isakova A, Deplancke B, Bernstein BE, et al. 2017. Systematic dissection of genomic features determining transcription factor binding and enhancer function. Proc Natl Acad Sci 114: E1291–E1300. http://www.pnas.org/lookup/doi/10.1073/pnas.1621150114.

Guillemot F, Hassan BA. 2017. Beyond proneural: emerging functions and regulations of proneural proteins. Curr Opin Neurobiol 42: 93–101. https://linkinghub.elsevier.com/retrieve/pii/S0959438816302392.

Haider NB, Jacobson SG, Cideciyan A V., Swiderski R, Streb LM, Searby C, Beck G, Hockey R, Hanna DB, Gorman S, et al. 2000. Mutation of a nuclear receptor gene, NR2E3, causes enhanced S cone syndrome, a disorder of retinal cell fate. Nat Genet.

Hobert O. 2016. A map of terminal regulators of neuronal identity in Caenorhabditis elegans. Wiley Interdiscip Rev Dev Biol 5: 474–498.https://onlinelibrary.wiley.com/doi/10.1002/wdev.233.

Hobert O. 2002. PCR fusion-based approach to create reporter Gene constructs for expression analysis in transgenic C. elegans. Biotechniques 32: 728–730.

Hobert O. 2008. Regulatory logic of neuronal diversity: Terminal selector genes and selector motifs. Proc Natl Acad Sci 105: 20067–20071. http://www.pnas.org/cgi/doi/10.1073/pnas.0806070105.

Hobert O, Kratsios P. 2019. Neuronal identity control by terminal selectors in worms, flies, and chordates. Curr Opin Neurobiol 56: 97–105. https://linkinghub.elsevier.com/retrieve/pii/S0959438818302605.

Huang X, Tian E, Xu Y, Zhang H. 2009. The C. elegans engrailed homolog ceh-16 regulates the self-renewal expansion division of stem cell-like seam cells. Dev Biol.

Hynes M, Stone DM, Dowd M, Pitts-Meek S, Goddard A, Gurney A, Rosenthal A. 1997. Control of cell pattern in the neural tube by the zinc finger transcription factor and oncogene Gli-1. Neuron 19: 15–26.

Kamath RS, Martinez-Campos M, Zipperlen P, Fraser AG, Ahringer J. 2001. Effectiveness of specific RNA-mediated interference through ingested double-stranded RNA in Caenorhabditis elegans. Genome Biol.

Kerk SY, Kratsios P, Hart M, Mourao R, Hobert O. 2017. Diversification of C. elegans Motor Neuron Identity via Selective Effector Gene Repression. Neuron 93: 80–98. http://dx.doi.org/10.1016/j.neuron.2016.11.036.

Kheradpour P, Ernst J, Melnikov A, Rogov P, Wang L, Zhang X, Alston J, Mikkelsen TS, Kellis M. 2013. Systematic dissection of regulatory motifs in 2000 predicted human enhancers using a massively parallel reporter assay. Genome Res 23: 800–811.

Kratsios P, Kerk SY, Catela C, Liang J, Vidal B, Bayer EA, Feng W, De La Cruz ED, Croci L, Giacomo Consalez G, et al. 2017. An intersectional gene regulatory strategy defines subclass diversity of C. Elegans motor neurons. Elife 6: 1–31.

Leyva-Díaz E, Masoudi N, Serrano-Saiz E, Glenwinkel L, Hobert O. 2020. Brn3/POU-IV-type POU homeobox genes—Paradigmatic regulators of neuronal identity across phylogeny. WIREs Dev Biol 9: 1–30. https://onlinelibrary.wiley.com/doi/10.1002/wdev.374.

Liu A, Niswander LA. 2005. Bone morphogenetic protein signalling and vertebrate nervous system development. Nat Rev Neurosci.

Lloret-Fernández C, Maicas M, Mora-Martínez C, Artacho A, Jimeno-Martín Á, Chirivella L, Weinberg P, Flames N. 2018. A transcription factor collective defines the HSN serotonergic neuron regulatory landscape. Elife 7. https://elifesciences.org/articles/32785.

Maglich JM, Sluder A, Guan X, Shi Y, McKee DD, Carrick K, Kamdar K, Willson TM, Moore JT. 2001. Comparison of complete nuclear receptor sets from the human, Caenorhabditis elegans and Drosophila genomes. Genome Biol.

Maicas M, Jimeno-Martín Á, Millán-Trejo A, Alkema MJ, Flames N. 2021. The transcription factor LAG-1/CSL plays a Notch-independent role in controlling terminal differentiation, fate maintenance, and plasticity of serotonergic chemosensory neurons. PLoS Biol 19: 1–25.

Mall M, Kareta MS, Chanda S, Ahlenius H, Perotti N, Zhou B, Grieder SD, Ge X, Drake S, Euong Ang C, et al. 2017. Myt1l safeguards neuronal identity by actively repressing many non-neuronal fates. Nature.

Masoudi N, Yemini E, Schnabel R, Hobert O. 2021. Piecemeal regulation of convergent neuronal lineages by bHLH transcription factors in Caenorhabditis elegans. Development 148: 1–15. https://journals.biologists.com/dev/article/148/11/dev199224/269057/Piecemeal-regulation-of-convergent-neuronal.

Masserdotti G, Gascón S, Götz M. 2016. Direct neuronal reprogramming: Learning from and for development. Dev.

Mears AJ, Kondo M, Swain PK, Takada Y, Bush RA, Saunders TL, Sieving PA, Swaroop A. 2001. Nrl is required for rod photoreceptor development. Nat Genet.

Narasimhan K, Lambert SA, Yang AWH, Riddell J, Mnaimneh S, Zheng H, Albu M, Najafabadi HS, Reece-Hoyes JS, Fuxman Bass JI, et al. 2015. Mapping and analysis of Caenorhabditis elegans transcription factor sequence specificities. Elife.

Nuez I, Félix MA. 2012. Evolution of susceptibility to ingested double-stranded rnas in Caenorhabditis nematodes. PLoS One.

Poole RJ, Bashllari E, Cochella L, Flowers EB, Hobert O. 2011. A Genome-Wide RNAi screen for factors involved in neuronal specification in Caenorhabditis elegans. PLoS Genet.

Reilly MB, Cros C, Varol E, Yemini E, Hobert O. 2020. Unique homeobox codes delineate all the neuron classes of C. elegans. Nature 584: 595–601. http://dx.doi.org/10.1038/s41586-020-2618-9.

Remesal L, Roger-Banyat I, Chirivella L, Maicas M, Brocal-Ruiz R, Pérez-Villalva A, Cucarella C, Casado M, Flames N. PBX1 is a terminal selector of olfactory bulb dopaminergic neurons. Dev.

Remesal L, Roger-Baynat I, Chirivella L, Maicas M, Brocal-Ruiz R, Pérez-Villalba A, Cucarella C, Casado M, Flames N. 2020. PBX1 acts as terminal selector for olfactory bulb dopaminergic neurons. Development 147: dev.186841. https://journals.biologists.com/dev/article/doi/10.1242/dev.186841/266880/PBX1-acts-as-terminal-selector-for-olfactory-bulb.

Rentzsch F, Layden M, Manuel M. 2017. The cellular and molecular basis of cnidarian neurogenesis. Wiley Interdiscip Rev Dev Biol 6: 1–19.

Roy K, Kuznicki K, Wu Q, Sun Z, Bock D, Schutz G, Vranich N, Monaghan AP. 2004. The tlx gene regulates the timing of neurogenesis in the cortex. J Neurosci.

Shimizu T, Nakazawa M, Kani S, Bae YK, Shimizu T, Kageyama R, Hibi M. 2010. Zinc finger genes Fezf1 and Fezf2 control neuronal differentiation by repressing Hes5 expression in the forebrain. Development 137: 1875–1885.

Shirasaki R, Pfaff SL. 2002. Transcriptional Codes and the Control of Neuronal Identity. Annu Rev Neurosci.

Simmer F, Moorman C, Van Der Linden AM, Kuijk E, Van Den Berghe PVE, Kamath RS, Fraser AG, Ahringer J, Plasterk RHA. 2003. Genome-wide RNAi of C. elegans using the hypersensitive rrf-3 strain reveals novel gene functions. PLoS Biol.

Spitz F, Furlong EEM. 2012. Transcription factors: From enhancer binding to developmental control. Nat Rev Genet.

Stefanakis N, Carrera I, Hobert O. 2015. Regulatory Logic of Pan-Neuronal Gene Expression in C. elegans. Neuron 87: 733–750. https://linkinghub.elsevier.com/retrieve/pii/S089662731500673X.

Stegmaier P, Kel AE, Wingender E. 2004. Systematic DNA-binding domain classification of transcription factors. Genome Inform.

Sural S, Hobert O. 2021. Nematode nuclear receptors as integrators of sensory information. Curr Biol 31: 4361–4366.e2. https://doi.org/10.1016/j.cub.2021.07.019.

Swoboda P, Adler HT, Thomas JH. 2000. The RFX-Type Transcription Factor DAF-19 Regulates Sensory Neuron Cilium Formation in C. elegans. Mol Cell 5: 411–421. https://linkinghub.elsevier.com/retrieve/pii/S1097276500804360.

Taubert S, Ward JD, Yamamoto KR. 2011. Nuclear hormone receptors in nematodes: Evolution and function. Mol Cell Endocrinol.

Taylor SR, Santpere G, Weinreb A, Barrett A, Reilly MB, Xu C, Varol E, Oikonomou P, Glenwinkel L, McWhirter R, et al. 2021. Molecular topography of an entire nervous system. Cell 184: 4329–4347.e23. https://linkinghub.elsevier.com/retrieve/pii/S0092867421007583.

Tewhey R, Kotliar D, Park DS, Liu B, Winnicki S, Reilly SK, Andersen KG, Mikkelsen TS, Lander ES, Schaffner SF, et al. 2016. Direct identification of hundreds of expression-modulating variants using a multiplexed reporter assay. Cell 165: 1519–1529. https://doi.org/10.1016/j.cell.2016.04.027.

Thor S, Andersson SGE, Tomlinson A, Thomas JB. 1999. A LIM-homeodomain combinatorial code for motor-neuron pathway selection system, although expression was also observed in precursors of the. Nature 397: 76–80.

Van Auken K, Weaver D, Robertson B, Sundaram M, Saldi T, Edgar L, Elling U, Lee M, Boese Q, Wood WB. 2002. Roles of the homethorax/Meis/Prep homolog UNC-62 and the Exd/Pbx homologs CEH-20 and CEH-40 in C. elegans embryogenesis. Development.

Waclaw RR, Allen ZJ, Bell SM, Erdélyi F, Szabó G, Potter SS, Campbell K. 2006. The zinc finger transcription factor Sp8 regulates the generation and diversity of olfactory bulb interneurons. Neuron 49: 503–516.

Wende H, Lechner SG, Cheret C, Bourane S, Kolanczyk ME, Pattyn A, Reuter K, Munier FL, Carroll P, Lewin GR, et al. 2012. The transcription factor c-Maf controls touch receptor development and function. Science (80-) 335: 1373–1376.

Zetterström RH, Solomin L, Jansson L, Hoffer BJ, Olson L, Perlmann T. 1997. Dopamine neuron agenesis in Nurr1-deficient mice. Science (80-).

Zhao C, Emmons SW. 1995. A transcription factor controlling development of peripheral sense organs in C. elegans. Nature 373: 74–78.

Zheng X, Chung S, Tanabe T, Sze JY. 2005. Cell-type specific regulation of serotonergic identity by the C. elegans LIM-homeodomain factor LIM-4. Dev Biol.

