## Supplemental Material for "Joint actions of diverse transcription factor families establish neuron-type identities and promote enhancer selectivity"

**Supplemental Material and Methods**

**Supplemental Figure S1:** *cat-1p::gfp* expression in controls (L4440 empty vector) and *gfp* RNAi treated *rrf-3(pk1426)* strain.

**Supplemental Figure S2:** TF family distribution in each neuron type.

**Supplemental Figure S3:** PDE lineage analysis in *unc-62, lin-14, vab-5* and *hbl-1* mutants.

**Supplemental Figure S4**: RNAi phenotypes associated to known dopaminergic terminal selectors.

**Supplemental Figure S5:** Additional reporter expression analysis in *unc-62* and *vab-3* mutants.

**Supplemental Figure S6**: Lineage analysis in *unc-62* and *vab-3* mutants.

**Supplemental Figure S7**: Cell autonomous rescue of *vab-3(ot346)* ADE neuron phenotype with VAB-3 isoform a and mouse PAX6.

**Supplemental Figure S8**: Dopaminergic regulatory signature overlaps with experimentally isolated *cis* regulatory modules driving dopaminergic effector gene expression.

**Supplemental Figure S9**: Distribution of dopaminergic regulatory signature with flexible composition of TFBS.

**Supplemental Figure S10**: Dopaminergic regulatory signature distribution in other gene classes expressed in dopaminergic neurons**.**

**Supplemental Table S1.** TF RNAi library information. (xls file)

**Supplemental Table S2.** TF RNAi screen results. (xls file)

**Supplemental Table S3.** Known regulators of MA identity. (xls file)

**Supplemental Table S4.** Mutant validation. (xls file)

**Supplemental Table S5.** *C. elegans* neuronal mutants. (xls file)

**Supplemental Table S6.** Mouse and *Drosophila* genomic and neuronal-mutant TFs. (xls file)

**Supplemental Table S7.** Gene lists for bioinformatics analysis and distribution of ETS+HD+PBX regulatory signature. (xls file)

**Supplemental Table S8.** Scoring's raw data and statistics. (xls file)

**Supplemental Table S9.** Summary of expression and phenotypes of dopaminergic terminal selector collective. (At the end of this pdf document)

**Supplemental Table S10.** Strains. (xls file)

**Supplemental Table S11.** Primers. (xls file)

**Supplemental Material and Methods**

**Generation of the Transcription factor RNAi library**

702 clones were extracted from the published genome libraries of Dr. Ahringer's library (BioScience) and Dr. Vidal's library (BioScience) (Kamath et al., 2003; Rual et al., 2004). Clones were verified by Sanger sequencing. 170 clones were newly generated (Supplemental Table S1). TF clones were generated from genomic N2 DNA by PCR amplification of the target gene, sub-cloned into L4440 plasmid (pPD129.36, Addgene) and transformed into *E. coli* HT115 (ED3) strain (from CGC). The complete list of TF RNAi feeding clones and primers used to generate new clones are listed in (Supplemental Table S1).

**RNAi feeding experiments and cell type specific RNAi**

For RNAi feeding experiments, 6 mM Isopropyl β-D-1-thiogalactopyranoside (IPTG) was added to NGM medium to prepare RNAi plates. RNAi clones were cultured overnight and induced with IPTG (4 mM) three hours before seeding. Adult gravid hermaphrodites were transferred to seeded IPTG plates within a drop of alkaline hypochlorite solution. After overnight incubation at 20 ºC, 10-15 newly hatched larvae were picked into fresh IPTG plates seeded with the same RNAi clone and considered the parental generation (P0). Approximately 7 days later young adult F1 generation was scored. Lethal RNAi clones, that precluded F1 analysis, were scored at P0 as young adults (Supplemental Table S2). In each replicate, a minimum of 30 worms per RNAi clone, coming from three distinct plates were scored. All experiments were performed at 20 °C. Each clone was scored in two independent replicates, a third replicate was performed when results from the first and second replicates did not coincide. In all experiments we included L4440 empty clone as negative control and *gfp* RNAi clone as positive control.

For cell type specific RNAi, 66 ng/µl of each sense and antisense PCR products were mixed together with *ttx-3p::mcherry* (33 ng/µl) and *rol-6(su1006)* (33 ng/µl) as co-injection markers. Primers used for this procedure are listed in (Supplemental Table S10).

**Generation of *C. elegans* alleles and transgenic lines**

Endogenous tagging of *cat-2(syb4698) and dat-1(syb4741)* was achieved by the insertion of a T2A::mNeonGreen fragment into the C-terminal region of the corresponding genes. The *syb5072* mutation was introduced in *cat-2(syb4698)* allele by point-mutation of a putative MEIS binding site from the wild type sequence GTGTC (located 105 base pairs upstream the ATG of the *cat-2* gene) to the mutated sequence CTTTA. The *syb5026* mutation was introduced in *cat-2(syb4698)* allele by the point-mutation of a putative PAIRED binding site from the wild type sequence GGAGCAAC (located 68 base pairs upstream the ATG of the *cat-2* gene) to the mutated sequence GGCTTAAC. All knock-in and point-mutated strains were verified by DNA sequencing. For the identification of putative binding sites in the *cis*-regulatory analysis the following consensus sequences were used: GACA for UNC-62/MEIS HD (Campbell and Walthall, 2016) and GRAGBA for VAB-3/PAIRED HD (Holst et al., 1997; Kim et al., 2008). For microinjections the plasmid of interest (50 ng/µl) was injected together with *rol-6* co-marker pRF4[*rol-6(su1006)*, 100 ng/µl]. For rescue experiments, cDNAs corresponding to the entire coding sequence of *vab-3* and Pax6 were amplified by PCR and cloned into *dat-1* promoter reporter plasmids replacing *gfp* cDNA (See primers and plasmids in Supplemental Table S10). cDNA plasmid (25 ng/µl) was injected directly into the corresponding mutant background as complex arrays together with digested *E. coli* genomic DNA (50 ng/µl) and pNF417(*unc-122::rfp,* 25 ng/µl) fluorescent co-marker. Reporters used in the analysis of the dopaminergic regulatory signature, were built by fusion PCR with an NLS version of the pPD95.75 plasmid was used (pNF400, see Supplemental Table S9). Transgenic DNA mix was composed of 50 ng/µl of the corresponding reporter fusion PCR plus 100 ng/µl of the pRF4 plasmid.

**Scoring and statistics**

For RNAi experiments, reporter analysis of the dopaminergic regulatory signature and *cis*-analysis, 30 young adult animals per line or per RNAi clone and replicate were analysed. For mutant analysis, 50 individuals were scored. RNAi experiments and reporter PDE lineage analyses were performed at 20 ºC while *cis*-regulatory and mutant analyses were performed at 25 ºC. For *cis*-regulatory analysis three independent lines for each transgenic construct were analysed.

Lack of GFP signalling was considered OFF; if GFP expression was substantially weaker than WT, a 'FAINT' category was included.

See Supplemental Table S8 for raw data on scorings, statistical tests and p values.

**Bioinformatics analysis**

Unless otherwise indicated, all analyses were performed using the software R (R Core Team, 2021) and packages from Bioconductor (Huber et al., 2015).

**Gene expression heatmap for monoaminergic neurons**

For generating the expression heatmap (Figure 1), scRNA-seq data from L4 *C. elegans larvae* (Taylor et al., 2021) was used. Only genes having more than 2 TPM in at least one cell type (a total of 13081 genes) were used to calculate the Z-score. Spearman rank correlation distance metrics of the Z-score matrix was used as base for hierarchical clustering of genes and samples using the complexHeatmap R package.

**TF family distribution in different species**

To infer the TF family distribution in *C. elegans*, *Drosophila* and mouse, we used data from WormBase (http://www.wormbase.org, version WS282, date of last access November of 2021), FlyBase (http://flybase.org/, version FB2020_01, date of last access April of 2021) and Mouse Genome Informatics (http://www.informatics.jax.org/, date of last access June of 2021).

To retrieve TFs with associated neuronal phenotypes in *C. elegans* we filtered all TFs with alleles associated to phenotypes (not RNAi evidence), and then manually curated these to get a list of neuronal-associated phenotypes, which were fount for only 93 TFs in *C. elegans* (out of 875 total TFs). A similar strategy was followed to retrieve TFs with associated neuronal phenotypes for *Drosophila* (266 out of 682 total TFs) and mouse (473 out of 1483 total TFs) TF alleles.

**DA signature analysis**

scRNA-seq data from Cao et al. (2017) was downloaded from the author's website (http://atlas.gs.washington.edu/worm-rna/). These data correspond to nearly 50,000 cells coming from 27 different cell types of the nematode *Caenorhabditis elegans* at the L2 larval stage. Cells with failed QC, doublets and unclassified cells were excluded from the subsequent analysis. Differential expression analysis between the cluster of dopaminergic neurons and the ciliated sensory neurons was performed using Monocle (Trapnell et al., 2014; Qiu et al. 2017a, 2017b) and results were filtered by q-value ($\leq$ 0.05) in order to get a list of differentially expressed genes in dopaminergic neurons. For clusters corresponding to non-DA neurons (RIA, ASE, Touch receptor neurons, GABAergic neurons and ALN/PLN/SDQ neurons) we followed a similar strategy, performing differential expression tests between each one of the selected neuronal clusters and all cells from the dataset annotated as neurons. Specificity of these six gene sets was checked by the enrichment of its anatomical association in *C. elegans* using the web tool WormEnrichr (https://amp.pharm.mssm.edu/WormEnrichr/) (Chen et al., 2013; Kuleshov et al., 2016). Gene lists for ubiquitous, panneuronal and panciliated categories were inferred from a more comprehensive scRNA-seq dataset (Packer et al.

2019). Briefly, we retrieved gene expression data (log2 transcripts per million) from all the genes that were expressed in at least one annotated terminal cell bin, getting a final matrix of 15,813 genes x 409 terminal cell bins. Following authors' original approach, genes were ordered by hierarchical clustering and cell bins were ordered by tissues; resulting in differential gene clusters which marked sites of predominant expression [Figure S32 from (Packer et al. 2019)]. Using these data, we manually curated three comprehensive gene lists: ubiquitous, panneuronal, and panciliated expression (Supplemental Table S7). For dopaminergic regulatory signature analysis, when genes were located in operons, only the gene located at the 5' end of the cluster, and thus subdued to *cis*-regulation, was considered, removing a total of 2,083 genes from the analysis. For hybrid operons, additional promoters were also included (Blumenthal, Davis & Garrido-Lecca, 2015). The final curated gene lists used in this study are listed in Supplemental Table S7.

For *C. elegans* regulatory signature analysis, we downloaded PWMs from CisBP version 1.02 (Weirauch et al. 2014) corresponding to the TF binding sites of the five transcription factors that compose the dopaminergic regulatory signature in *C. elegans*. If the exact match for *C. elegans* was not available, we selected the PWM from the *M. musculus* or *H. sapiens* orthologous TFs (ETS, ref. M0709; HD, ref. M5340; PBX, ref. M1898; MEIS, ref. M6048; PAIRED, ref. M1500), plus an additional hybrid PAIRED HD site (represented as HD*, ref. M6189). Following published methodology (Lloret-Fernández et al. 2018) we downloaded upstream and intronic gene regions of protein-coding genes from WormBase (version 262) and then classified genes using the gene lists mentioned above. Upstream regions were trimmed to a maximum of 10 kb. PWMs were aligned to genomic sequences and we retrieved matches with a minimum score of 70%. To increase specificity, we removed all matches that did not bear an exact consensus sequence for the corresponding TF family (consensus sequence for ETS: VMGGAWR, HD: TAATT, PBX: GATNNAT, MEIS: DTGTCD, HD*: HTAATTR, PAIRED: GGAAC). Sliding window search with a maximum length of 700 bp was performed to find regions that included at least one match for the 6 TF binding motifs, allowing flexible motif composition. To assess signature enrichment in the set of dopaminergic expressed genes, 10,000 sets of 86 random genes were built considering that: 1) they were not differentially expressed in dopaminergic neurons, 2) at least one ortholog had been described in other *Caenorhabditis* species (*C. brenneri, C. briggsae, C. japonica* or *C. remanei*), and 3) their upstream and intronic regions were similar in length, on average, to those of the dopamine-expressed genes (Mann-Whitney U test, p-value > 0.05). For each one of the non-dopaminergic neuronal groups (RIA, ASE, Touch receptor neurons,

GABAergic neurons and ALN/PLN/SDQ neurons), similar sets of random genes were built. We considered the enrichment in signature to be significant when the percentile of the neuronal group in regard to the internal random control was above 95. Differences between dopaminergic expressed genes and other neuronal groups were assessed by Brunner-Munzel test, performed with R package brunnermunzel (Toshiaki 2019), *: $p$-value < 0.05, **: $p$-value < 0.01, ***: $p$-value < 0.001).

**References**


Blumenthal, T., Davis, P., & Garrido-Lecca, A. (2015). Operon and non-operon gene clusters in the C. elegans genome. *WormBook : The Online Review of C. Elegans Biology*. https://doi.org/10.1895/wormbook.1.175.1

Cao, J., Packer, J. S., Ramani, V., Cusanovich, D. A., Huynh, C., Daza, R., … Shendure, J. (2017). Comprehensive single-cell transcriptional profiling of a multicellular organism. *Science*, *357*(6352), 661–667. https://doi.org/10.1126/science.aam8940

Chen, E. Y., Tan, C. M., Kou, Y., Duan, Q., Wang, Z., Meirelles, G. V., ... Ma'ayan, A. (2013). Enrichr: Interactive and collaborative HTML5 gene list enrichment analysis tool. BMC Bioinformatics, 14. https://doi.org/10.1186/1471-2105-14-128

Eden, E., Navon, R., Steinfeld, I., Lipson, D., & Yakhini, Z. (2009). GOrilla: A tool for discovery and visualization of enriched GO terms in ranked gene lists. *BMC Bioinformatics*, *10*. https://doi.org/10.1186/1471-2105-10-48

Huber, W., Carey, V. J., Gentleman, R., Anders, S., Carlson, M., Carvalho, B. S., … Morgan, M. (2015). Orchestrating high-throughput genomic analysis with Bioconductor. *Nature Methods*, *12*(2), 115–121. https://doi.org/10.1038/nmeth.3252

Kamath RS, Fraser AG, Dong Y, Poulin G, Durbin R, Gotta M, Kanapin A, Le Bot N, Moreno S, Sohrmann M, et al. 2003. Systematic functional analysis of the Caenorhabditis elegans genome using RNAi. *Nature*.

Kuleshov, M. V., Jones, M. R., Rouillard, A. D., Fernandez, N. F., Duan, Q., Wang, Z., ... Ma'ayan, A. (2016). Enrichr: a comprehensive gene set enrichment analysis web server 2016 update. Nucleic Acids Research, 44(W1), W90-7. https://doi.org/10.1093/nar/gkw377

Lloret-Fernández, C., Maicas, M., Mora-Martínez, C., Artacho, A., Jimeno-Martín, Á., Chirivella, L., … Flames, N. (2018). A transcription factor collective defines the HSN serotonergic neuron regulatory landscape. *ELife*, *7*. https://doi.org/10.7554/eLife.32785

Qiu, X., Mao, Q., Tang, Y., Wang, L., Chawla, R., Pliner, H., & Trapnell, C. (2017a). Reversed graph embedding resolves complex single-cell developmental trajectories. BioRxiv.

Qiu, X., Hill, A., Packer, J., Lin, D., Ma, Y. A., & Trapnell, C. (2017b). Single-cell mRNA quantification and differential analysis with Census. Nature Methods, 14(3), 309–315. https://doi.org/10.1038/nmeth.4150

R Core Team (2021). R: A language and environment for statistical computing. R Foundation for Statistical Computing, Vienna, Austria. URL: https://www.R-project.org/

Rual JF, Ceron J, Koreth J, Hao T, Nicot AS, Hirozane-Kishikawa T, Vandenhaute J, Orkin SH, Hill DE, van den Heuvel S, et al. 2004. Toward improving Caenorhabditis elegans phenome mapping with an ORFeome-based RNAi library. *Genome Res*.

Taylor SR, Santpere G, Weinreb A, Barrett A, Reilly MB, Xu C, Varol E, Oikonomou P, Glenwinkel L, McWhirter R, et al. 2021. Molecular topography of an entire nervous system. *Cell* 184: 4329-4347.e23.

Toshiaki Ara (2019). brunnermunzel: (Permuted) Brunner-Munzel Test. R package version 1.3.5. https://CRAN.R-project.org/package=brunnermunzel

Trapnell, C., Cacchiarelli, D., Grimsby, J., Pokharel, P., Li, S., Morse, M., ... Rinn, J. L. (2014). The dynamics and regulators of cell fate decisions are revealed by pseudotemporal ordering of single cells. Nature Biotechnology, 32(4), 381–386. https://doi.org/10.1038/nbt.2859

Van Rossum, G., & Drake Jr, F. L. (1995). *Python tutorial*. Centrum voor Wiskunde en Informatica Amsterdam, The Netherlands.

Weirauch, M. T., Yang, A., Albu, M., Cote, A. G., Montenegro-Montero, A., Drewe, P., ... Hughes, T. R. (2014). Determination and Inference of Eukaryotic Transcription Factor Sequence Specificity. Cell, 158(6), 1431–1443. https://doi.org/10.1016/j.cell.2014.08.009

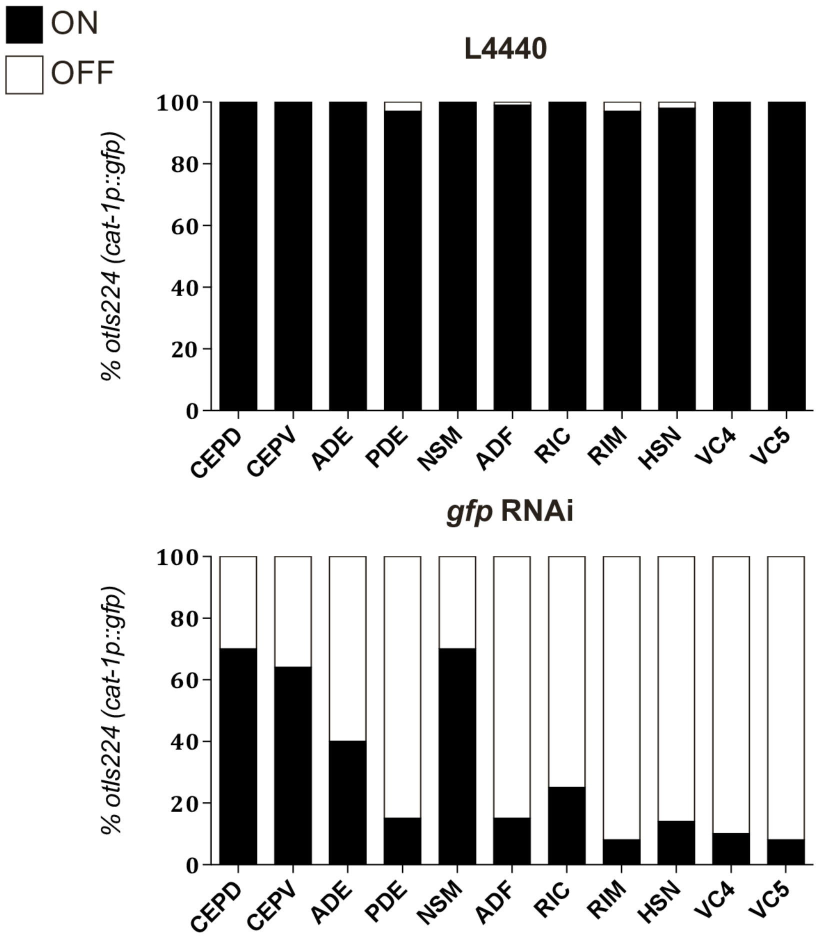


**Supplemental Figure 1. Analysis of *cat-1p::gfp* expression in controls (L4440 empty vector) and *gfp* RNAi treated *rrf-3(pk1426)* worms**. NSM and CEPs seem to be more refractory to RNAi effects than other MA neurons. n=30 worms per condition.

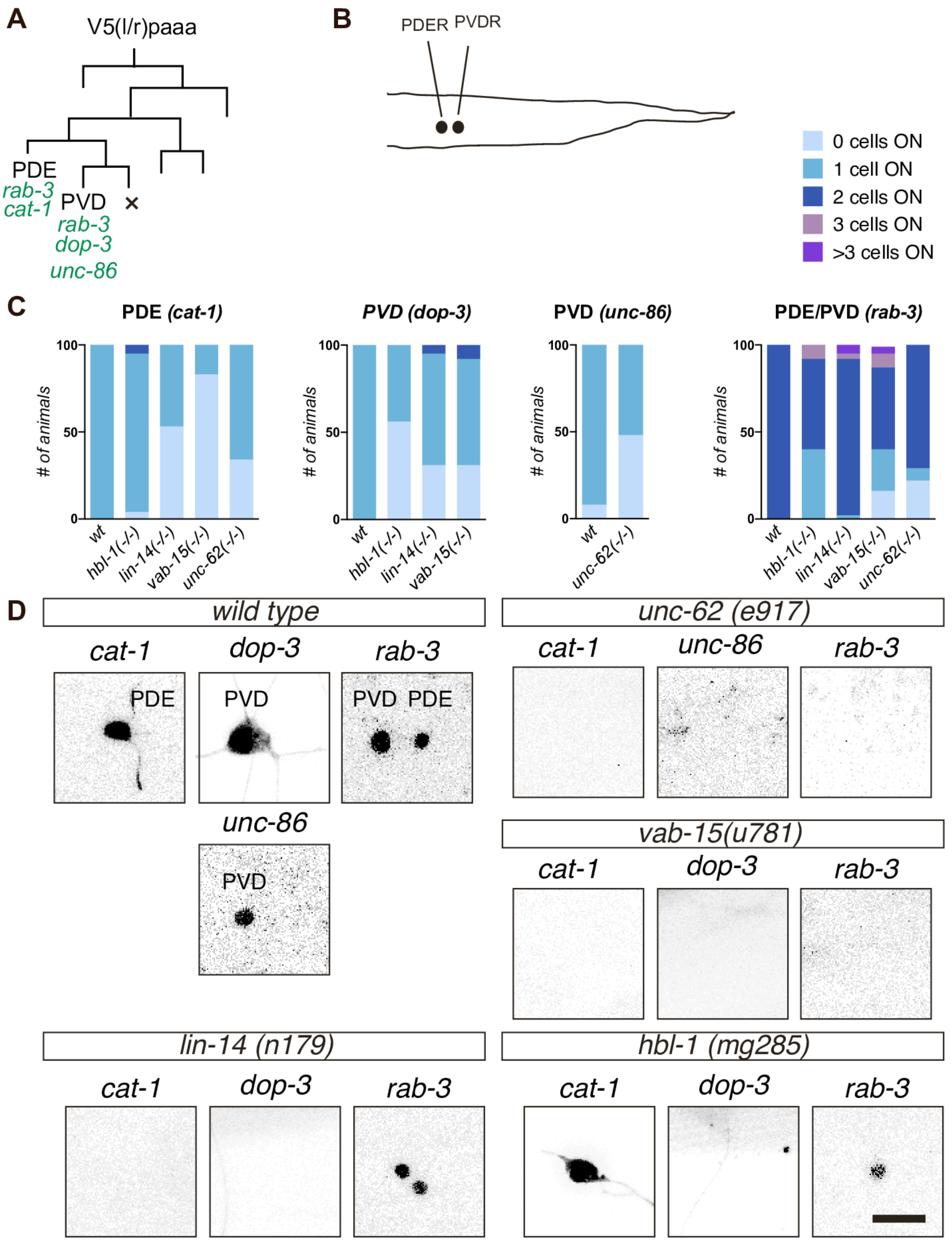


**Supplemental Figure 2. Analysis of PDE and PVD markers in *hbl-1, lin-14, vab-14* and *unc-62* TF mutants.**

**A**) V5 lineage representation. One of the lineage branches generates the PDE and PVD neuron and a cell death event. *rab-3* panneuronal reporter is expressed in both neurons, *cat-1* reporter is exclusively expressed in PDE while *unc-86* POU HD TF and *dop-3* dopamine receptor reporters are expressed in PVD.

**B**) PDER and PVDR neurons are located in the right side of the worm isolated from other neurons, allowing for easy identification of panneuronal reporter cells.

**C**) Quantification of *cat-1p::gfp(otIs221), dop-3p::rfp(vsIs33), unc-86::yfp(otIs337) and rab-3p::rfp(otIs355) or rab-3p::yfp (otIs287)* in *wild type, unc-62(e917), vab-15(u781), lin-14(n179)* and *hbl-1(mg285)* mutants. n>50 animals per genotype and reporter. See Supplemental Table S8 for raw data.

**D**) Representative micrographs of the corresponding mutant allele phenotypes. Scale: 10 µm.

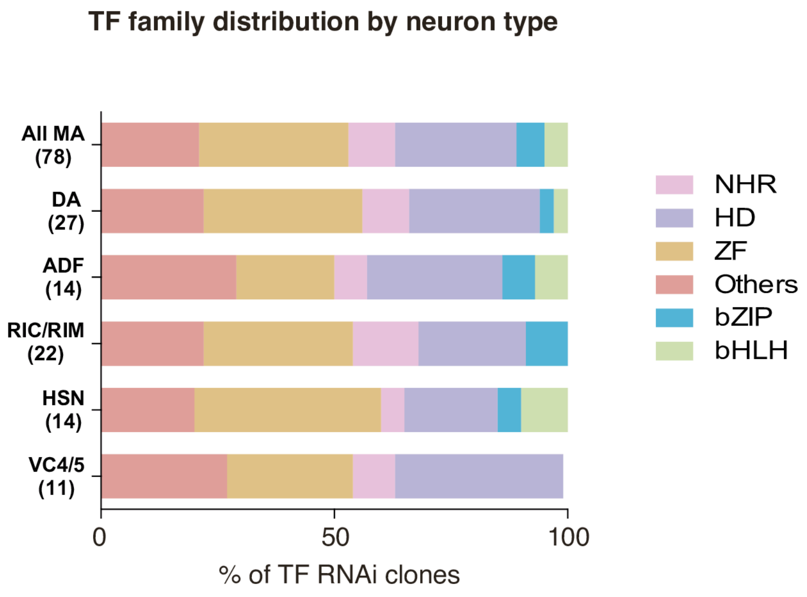


**Supplemental Figure 3. TF family distribution for TFs affecting reporter expression in each neuron type.** NSM is not included as only two RNAi clones produce NSM differentiation defects and the four dopaminergic neurons types are represented together as "DA" as they share many TFs RNAi clones affecting reporter expression.

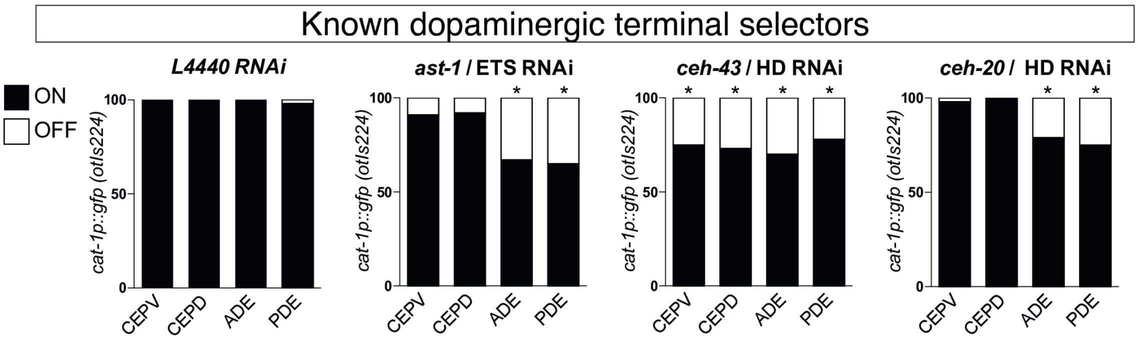


**Supplemental Figure 4. RNAi phenotypes associated to known dopaminergic terminal selectors.** RNAi against *ast-1, ceh-43* and *ceh-20,* known dopaminergic terminal selectors, produces significant GFP expression defects in at least two out of the four dopaminergic neuron types. *ceh-20* RNAi treated animals were scored at P0 due to embryonic lethal effects at F1. *: $p<0.05$. Fisher's exact test. n= 30 animals, at least two independent replicates per RNAi clone.

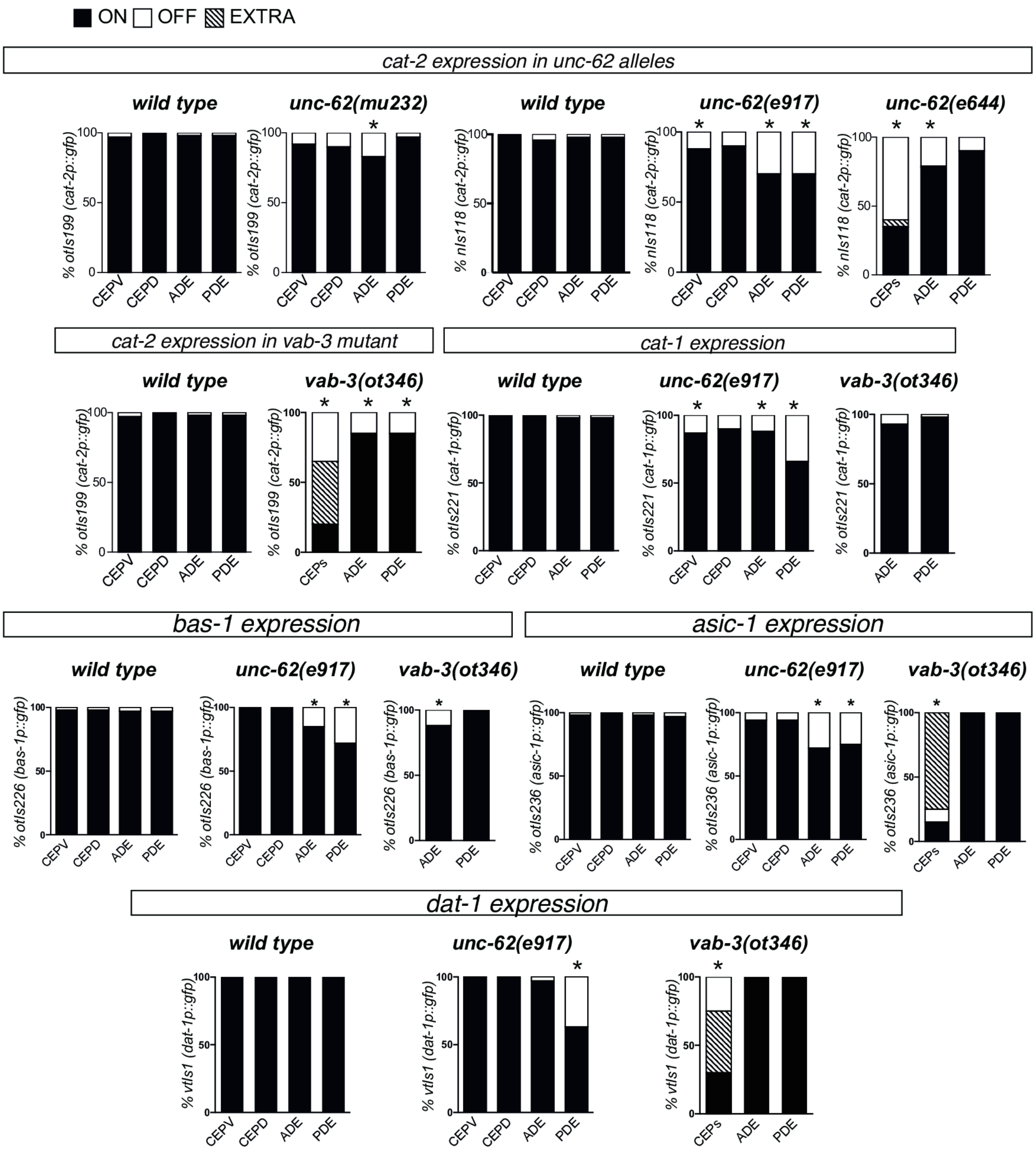


**Supplemental Figure 5. Transcriptional reporter expression analysis in *unc-62* and *vab-3* mutants.** For *cat-2*, *dat-1* and *asic-1* reporter analysis, disorganization of *vab-3(ot346)* head neurons precluded us from distinguishing CEPV from CEPD and thus are scored as a unique CEP category. For *bas-1* and *cat-1* reporter analysis disorganization of *vab-3(ot346)* head precluded the identification of CEPs among other GFP expressing neurons in the region and thus only ADE and PDE scorings are shown. n>50 animals each condition, *: p<0.05 compared to wild type.

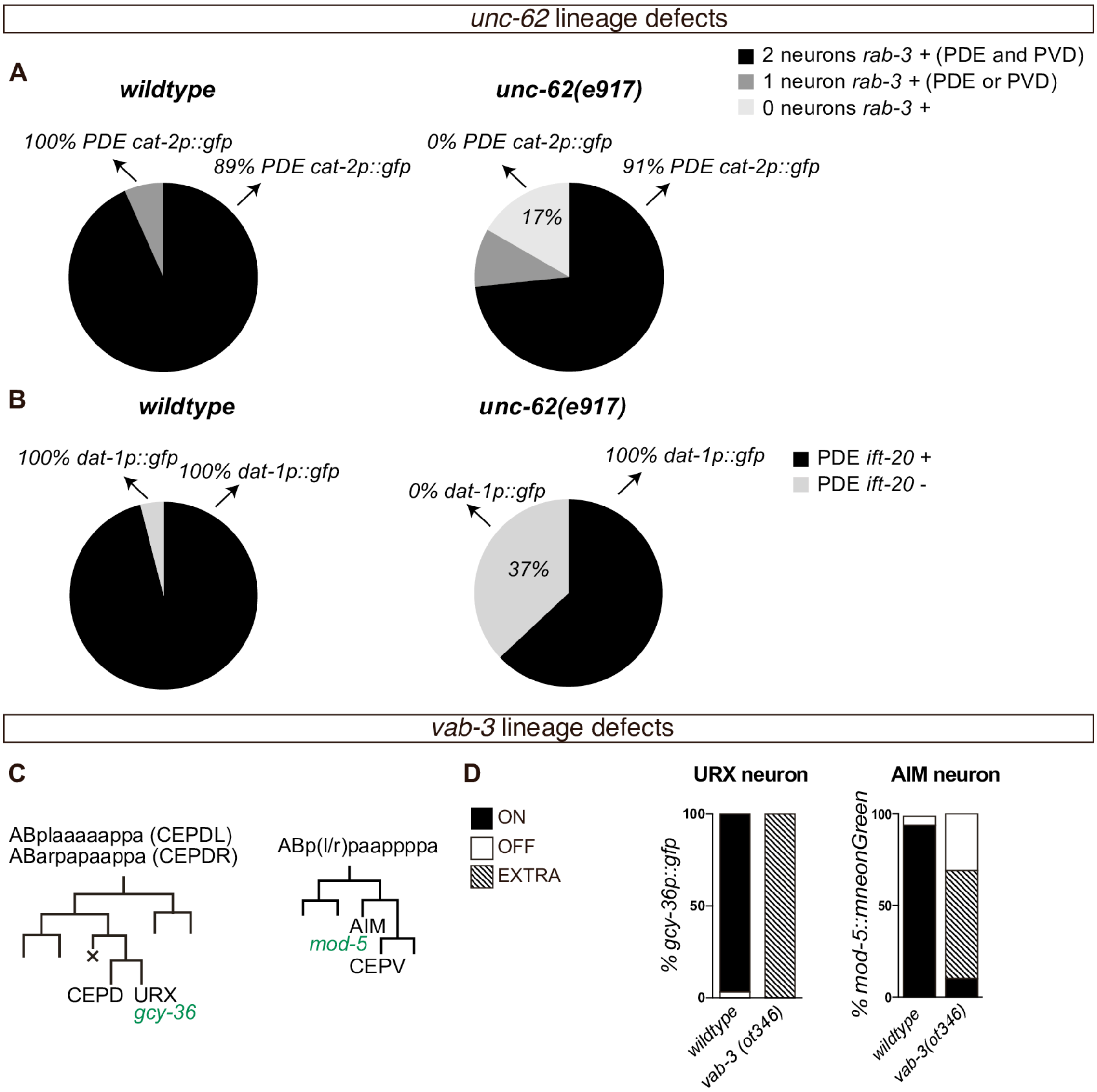


**Supplemental Figure 6. Lineage analysis in *unc-62* and *vab-3* mutants.** **A**) Co-expression analysis in PDE lineage of the panneuronal marker *rab-3* and dopaminergic marker *cat-2* in *unc-62(e917)* mutants. Most animals with *cat-2* expression defects in PDE also lack *rab-3* expression. n>30 scored neurons.

**B**) Co-expression analysis in *unc-62(e917)* PDE lineage of the panciliated marker *ift-20* and dopaminergic marker *dat-1*. Animals with missing *dat-1* expression in PDE also lack *ift-20* expression. n>30 scored neurons.

**C**) CEPV and CEPD lineage, in green genes expressed by the corresponding neuron.

**D**) *vab-3(ot346)* mutants show ectopic expression of *gcy-36* and *mod-5* confirming lineage defects.

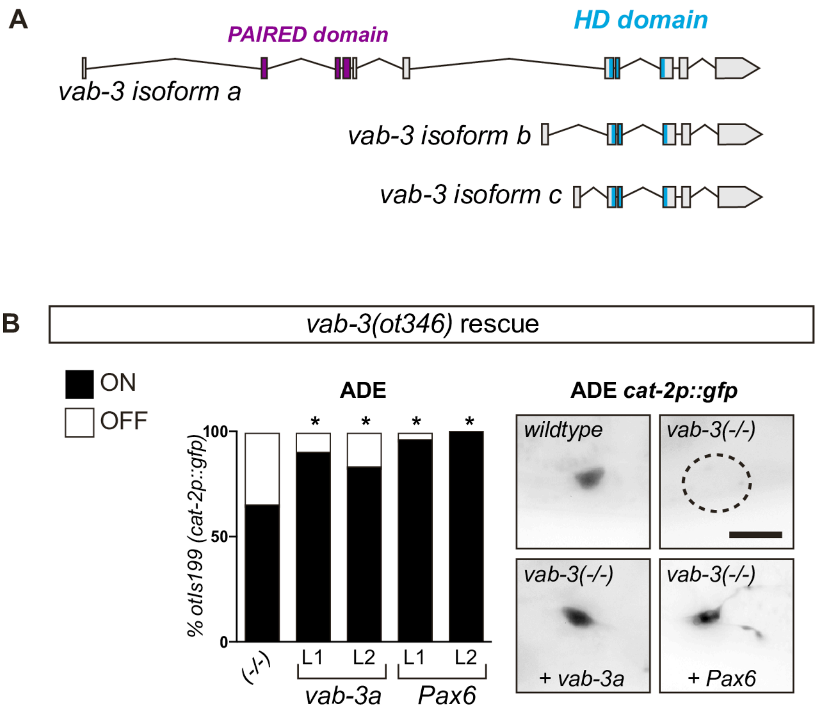


**Supplemental Figure 7. Cell autonomous rescue of *vab-3(ot346)* ADE neuron phenotype with VAB-3 isoform a and mouse PAX6.**

**A**) Schema of *vab-3* different coding isoforms. Only VAB-3a contains both the PAIRED and the HD DNA binding domains.

**B**) Quantification of *vab-3(ot346)* rescue experiments. *dat-1* promoter was used to drive specific dopaminergic expression of *vab-3 isoform a* cDNA and *Pax6* cDNA in *vab-3* mutants. ADE rescue defects of *cat-2* reporter expression was quantified as *dat-1* expression in this neuron is unaffected in *vab-3(ot346)* (Figure 2). Representative pictures of each experimental condition are shown in the right. L1 and L2 represent two independent transgenic lines. Scale: 10 µm. n>50 animals each condition, *: $p<0.05$ compared to mutant phenotype. Fisher's exact test with Bonferroni correction for multiple comparisons.

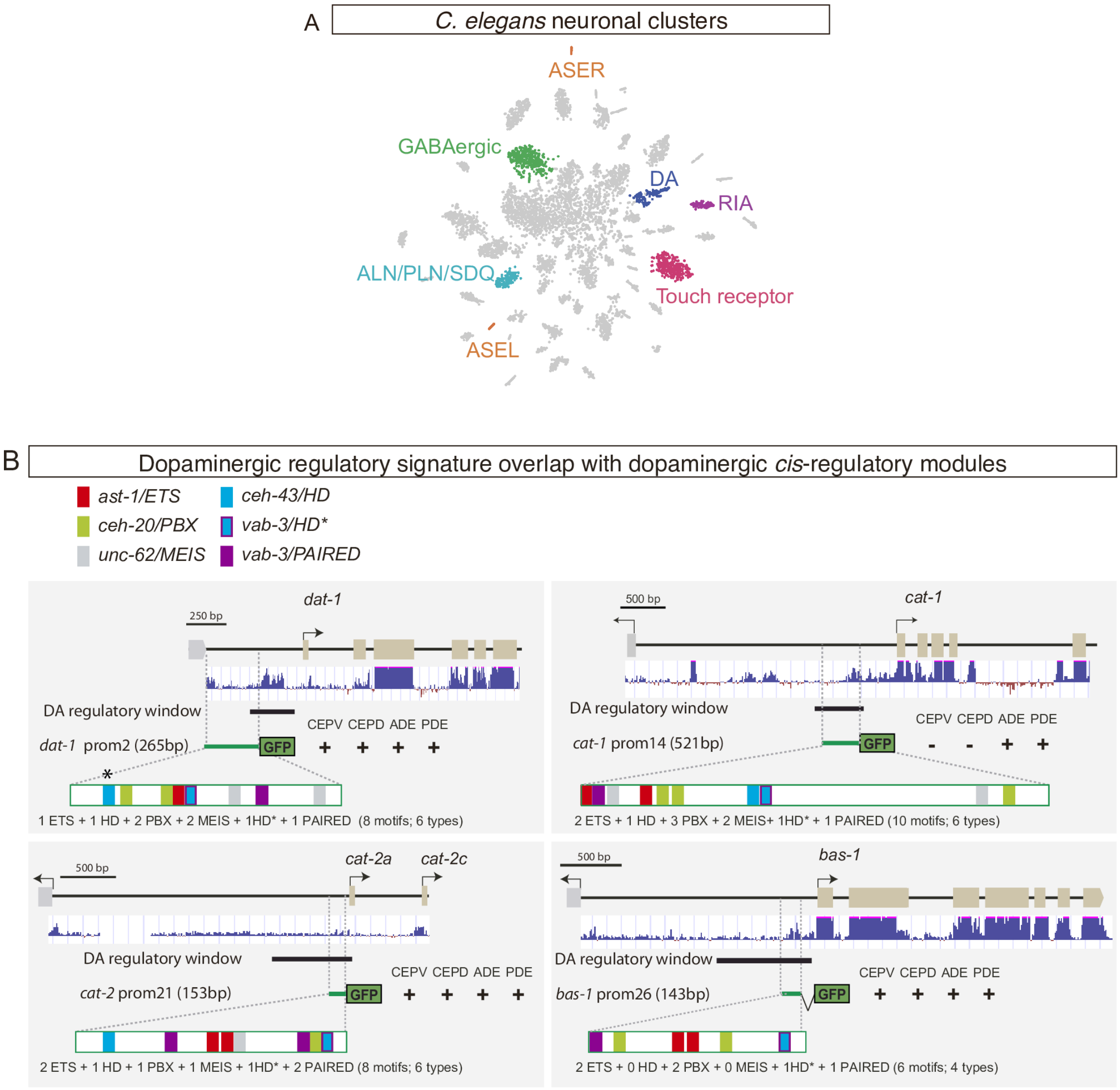


**Supplemental Figure 8. Dopaminergic regulatory signature overlaps with experimentally isolated *cis* regulatory modules driving dopaminergic effector gene expression.**

**A**) t-SNE visualization of high-level neuronal subtypes from single-cell analysis of L2 animals. Data obtained from (Cao et al., 2017). Neuronal types used in the dopaminergic regulatory signature analysis are highlighted in different colours.

**B**) Experimentally isolated minimal enhancers for four out of the five dopamine pathway genes (Flames and Hobert, 2009) overlap with predicted dopaminergic regulatory signature windows. Black lines represent the coordinates covered by the dopaminergic regulatory signature windows. Green lines mark published minimal enhancers for the respective gene (Doitsidou et al., 2013; Flames and Hobert, 2009). Specific matches for all six TFBS classes are represented with the color code indicated. In the *dat-1* gene, HD site marked with an asterisk indicates an experimentally identified site not retrieved in the bioinformatic analysis due to a small sequence mismatch. Dark blue bar profiles represent sequence conservation in *C. briggsae, C. brenneri, C. remanei* and *C. japonica*, showing that dopaminergic regulatory signature does not necessarily coincide with conserved regions.

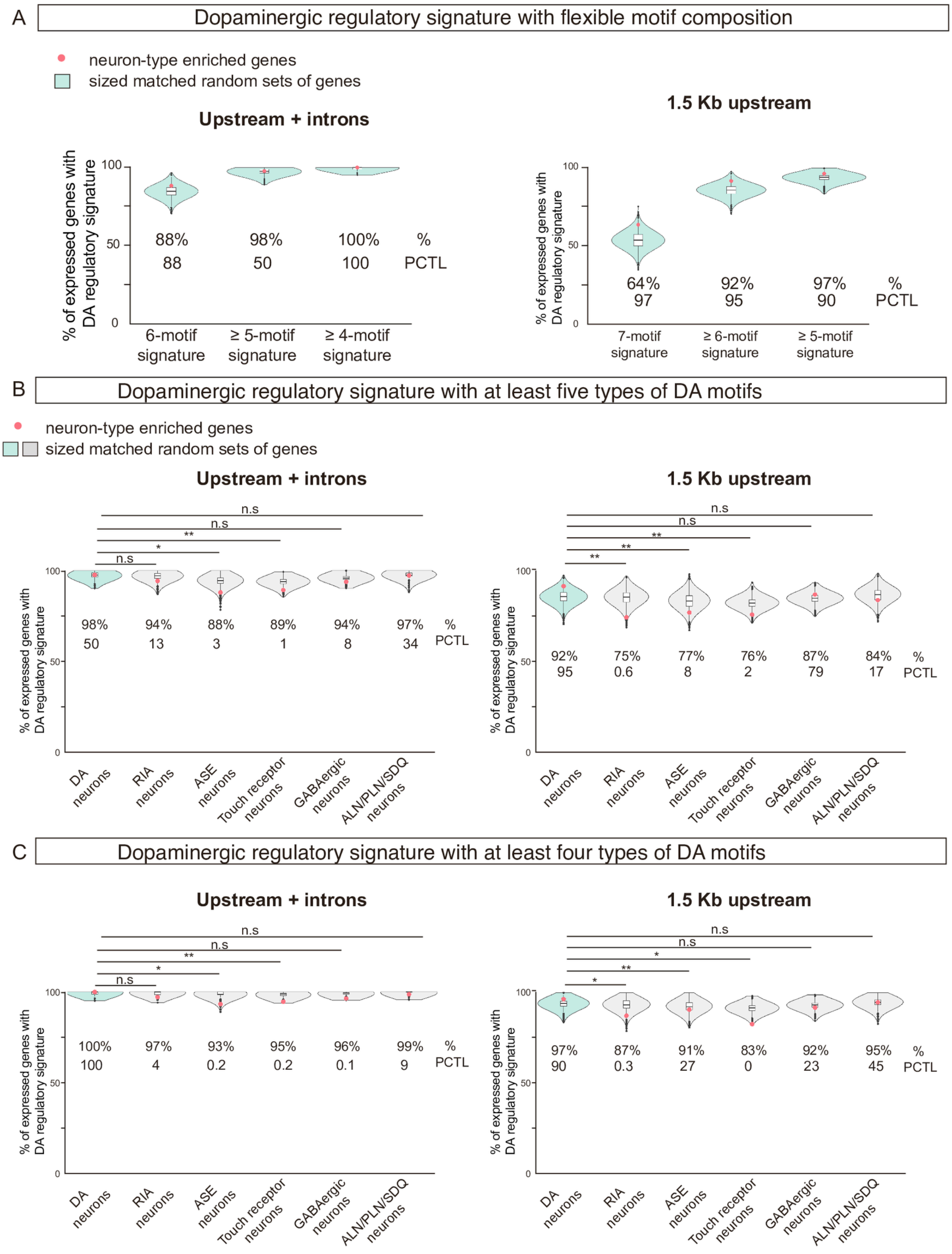

**Supplemental Figure 9. Distribution of dopaminergic degulatory signature with flexible composition of TFBS.**

**A**) Considering dopaminergic regulatory signature windows with at least five or at least four dopaminergic TF motifs increases the percentage of dopaminergic enriched genes with dopaminergic signature but concomitantly decreases enrichment compared to the 10,000 sets of random genes. %: percentage of genes with assigned dopaminergic regulatory signature. PCTL: percentile of the real value (red dot) in the 10,000 random set value distribution.

**B-C**) Analysis of windows with at least five or at least four motifs still shows higher percentage in dopaminergic expressed genes compared to some neuronal populations, particularly when considering only proximal regulatory regions (1.5 kb from ATG), however this difference is less pronounced than when considering the dopaminergic regulatory signature with the presence all six TFBS types (Figure 5). Brunner-Munzel test, n.s.: p>0.05.

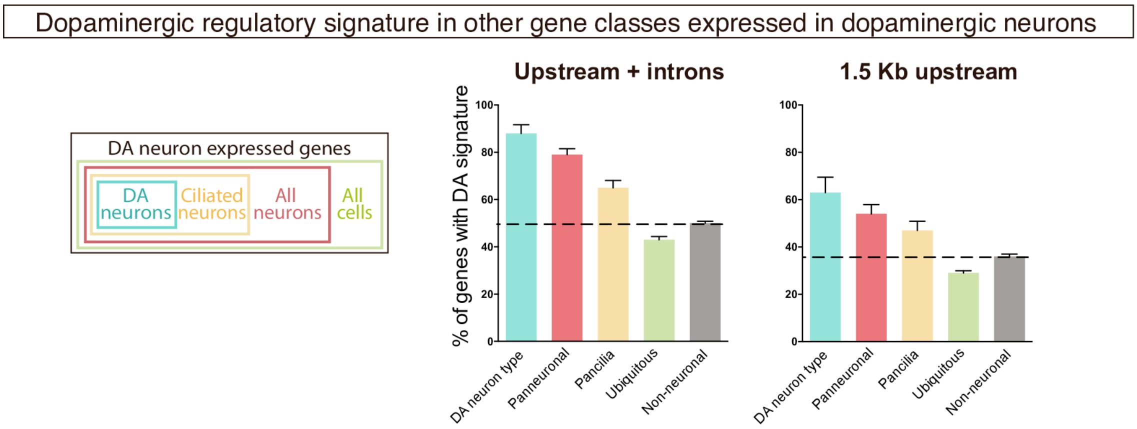


**Supplemental Figure 10. Dopaminergic regulatory signature distribution in other gene classes expressed in dopaminergic neurons.**

Four sets of genes associated to different patterns of gene expression specificity coexist in the dopaminergic neurons: 1) dopaminergic effector genes mostly specific of dopaminergic neurons and sometimes also in a few other neurons; 2) pancilia genes expressed by all sensory ciliated neurons; 3) panneuronal genes expressed by all neurons and 4) ubiquitous genes expressed by all cells. Dopaminergic regulatory signature is enriched in dopaminergic effector genes and to a less extent in panneuronal expressed and panciliated genes but is not present in ubiquitous genes. Quantification of dopaminergic signature in non-neuronal genes is shown as negative control and marked with a dashed line.

**Supplemental table S9**

**Summary of mutant phenotypes, *cis*-mutation effects and expression for the dopaminergic terminal selector collective.**

| | CEPV | CEPD | ADE | PDE |
|---|---|---|---|---|
| *ast-1* mutant phenotype | Yes | Yes | Yes | Yes |
| *ast-1* expression | Yes | Yes | Yes | Yes |
| ETS/*ast-1 cis* mutation effect | Yes | Yes | Yes | Yes |
| *ceh-43* mutant phenotype | Yes | Yes | Yes | Yes |
| *ceh-43* expression | Yes | Yes | Yes | Yes |
| HD/*ceh-43 cis* mutation effect | Yes | Yes | Yes | Yes |
| *ceh-20* mutant phenotype | No (*dat-1* reporter) | No (*dat-1* reporter) | No (*dat-1* reporter) | Yes (*dat-1* reporter) |
| *ceh-20* expression | Yes | Yes | Yes | Yes |
| PBX/*ceh-20 cis* mutation effect | Yes | Yes | Yes | Yes |
| *unc-62* mutant phenotype | Yes | Yes | Yes | Yes |
| *unc-62* expression | No | No | Yes | Yes |
| MEIS/ *unc-62 cis* mutation effect | Yes (Endogenous *cat-2* locus) | Yes (Endogenous *cat-2* locus and Ex array) | Yes (Endogenous *cat-2* locus and Ex array) | Yes (Ex array *cat-2*) |
| *vab-3* mutant phenotype | Yes | Yes | Yes | Yes |
| *vab-3* expression | Yes | Yes | No | No |
| PAIRED/vab-3 *cis* mutation effect | Yes (Endogenous *cat-2* locus) | Yes (Endogenous *cat-2* locus and Ex array) | Yes (Endogenous *cat-2* locus and Ex array) | Yes (Ex array *cat-2*) |

Source: This work and (Flames and Hobert 2009; Doitsidou et al. 2013; Reilly et al. 2020)
